# Astrocytes encode complex behaviorally relevant information

**DOI:** 10.1101/2021.10.09.463784

**Authors:** Katharina Merten, Robert W. Folk, Daniela Duarte, Axel Nimmerjahn

## Abstract

Astrocytes, glial cells of the central nervous system, help to regulate neural circuit operation and adaptation. They exhibit complex forms of chemical excitation, most prominently calcium transients, evoked by neuromodulator and -transmitter receptor activation^1–4^. However, whether and how astrocytes contribute to cortical processing of complex behavior remains unknown^1^. One of the puzzling features of astrocyte calcium transients is the high degree of variability in their spatial and temporal patterns under behaving conditions. Here, we provide mechanistic links between astrocytes’ activity patterns, molecular signaling, and behavioral cognitive and motor activity variables by employing a visual detection task that allows for in vivo calcium imaging, robust statistical analyses, and machine learning approaches. We show that trial type and performance levels deterministically shape astrocytes’ spatial and temporal response properties. Astrocytes encode the animals’ decision, reward, and sensory properties. Our error analysis confirms that astrocytes carry behaviorally relevant information depending on and complementing neuronal coding. We also report that cell-intrinsic mechanisms curb astrocyte calcium activity. Additionally, we show that motor activity-related parameters strongly impact astrocyte responses and must be considered in sensorimotor study designs. Our data inform and constrain current models of astrocytes’ contribution to complex behavior and brain computation beyond their established homeostatic and metabolic roles.

## Introduction

Mounting evidence from multiple species and central nervous system regions suggests that astrocytes play pivotal roles in neural circuit function and behavior^2, 3, 5^. Although not electrically excitable, astrocytes display a complex repertoire of intracellular signaling, most prominently calcium transients, triggered by neurotransmitter and neuromodulator receptor activation on their surface. This signaling spans multiple spatial and temporal scales, from sub-second transients in single astrocytes to seconds- or even minutes-long transients in astrocytic networks, suggesting that astrocytes may carry out computations on various timescales related to sensory processing, brain state modulation, and memory formation^1, 2, 4^.

Despite the recent technical progress in measuring neuronal, astrocyte, and transmitter dynamics in behaving animals, a key unresolved question is precisely how astrocyte excitation relates to animal behavior and how it may contribute to brain computation of cognitive functions. This knowledge gap is partly due to the lack of standardized quantitative behavioral assays that allow tight control over the animal’s behavior and associated cellular and molecular signaling.

Additionally, data analysis approaches are often based on manually drawn regions of interest, which are poorly suited to capture the complexity of astrocyte excitation or its relationship to circuit dynamics and behavior^6, 7^. Moreover, recent studies reporting astrocytic encoding of spatial information^8^ or reward location^9^ in the hippocampus have neglected the impact of mouse motor behavior on astrocyte responses. Astrocytes are known to exhibit widespread calcium excitation during locomotion mediated by local neurotransmitter and volumetric neuromodulator release^10–12^. Therefore, run parameters, particularly the timing of astrocyte response onset relative to run onset and the period between runs, might strongly influence experimental results and interpretation.

Using a quantitative visual detection task, in vivo calcium imaging, robust statistical analyses that account for the joint influence of run and cognitive parameters, and machine learning approaches, we show that astrocyte population transients (“syncytium responses”) in the mouse motor cortex are deterministic and encode information about the stimulus, trial type, reward, decision, and the animal’s performance level. Astrocyte responses were also significantly linked to run onset, run duration, and inter-run interval. Additionally, we show that astrocyte population responses underlie intrinsic constraints. Our data provide insight into fundamental computations within astrocyte networks and the integration and transformation of molecular signals within their environment, suggesting that these cells contribute to complex behavior and brain computation beyond their established homeostatic and metabolic roles.

## Results

To investigate whether astrocytes contribute to cortical information processing and the encoding of complex behavior, we recorded their activity in a visual detection task. This behavioral assay involved numerous trial repetitions across multiple sessions and allowed robust regression and decoding analysis. In total, we recorded 4,837 trials across 21 behavioral sessions (see **Methods**). Mice were trained to report the presence or absence of a visual stimulus by running on a spherical treadmill for a fluid reward (**Fig. 1a,b**). Stimuli were presented at two intensity levels: i) salient and ii) close to the animal’s perceptual level. The internal state of the mouse determined whether it had seen the stimulus (‘yes’ decision if the animal initiated a run during stimulus presentation; ‘no’ decision if it stood still for >3 s). Decision outcomes were classified according to signal detection theory (**Fig. 1c**). Before stimulus presentation, the mice were required to stand still for 20 s. If mice interrupted the stand-still phase, the trial was aborted and counted as a spontaneous run. **Fig. 1d** shows trial outcome proportions for an example session. To create psychometric detection curves for each animal, we used the proportion of ‘yes’ decisions for the two stimulus intensities (**Fig. 1e**). The mice were rewarded on correct trials only (hits and correct rejections). Trials with stimuli close to the perceptual threshold, in which the animals did not detect the stimulus, were not reinforced. This reward contingency led to a slight bias of the mice to erroneously report the presence of a stimulus in some of the stimulus absent trials (**Fig. 1e**). The animals’ performance levels varied within and across sessions. We computed a measure of discriminability (d’) derived from signal detection theory^13^ by subtracting z-scores (normal deviates) of median ‘hit’ rates from z-scores of median ‘false alarm’ rates (**Fig. 1f**). Performance intervals exceeding the detection threshold were considered high- performance states, whereas periods with d’<2 were classified as low-performance states.

**Fig. 1.**
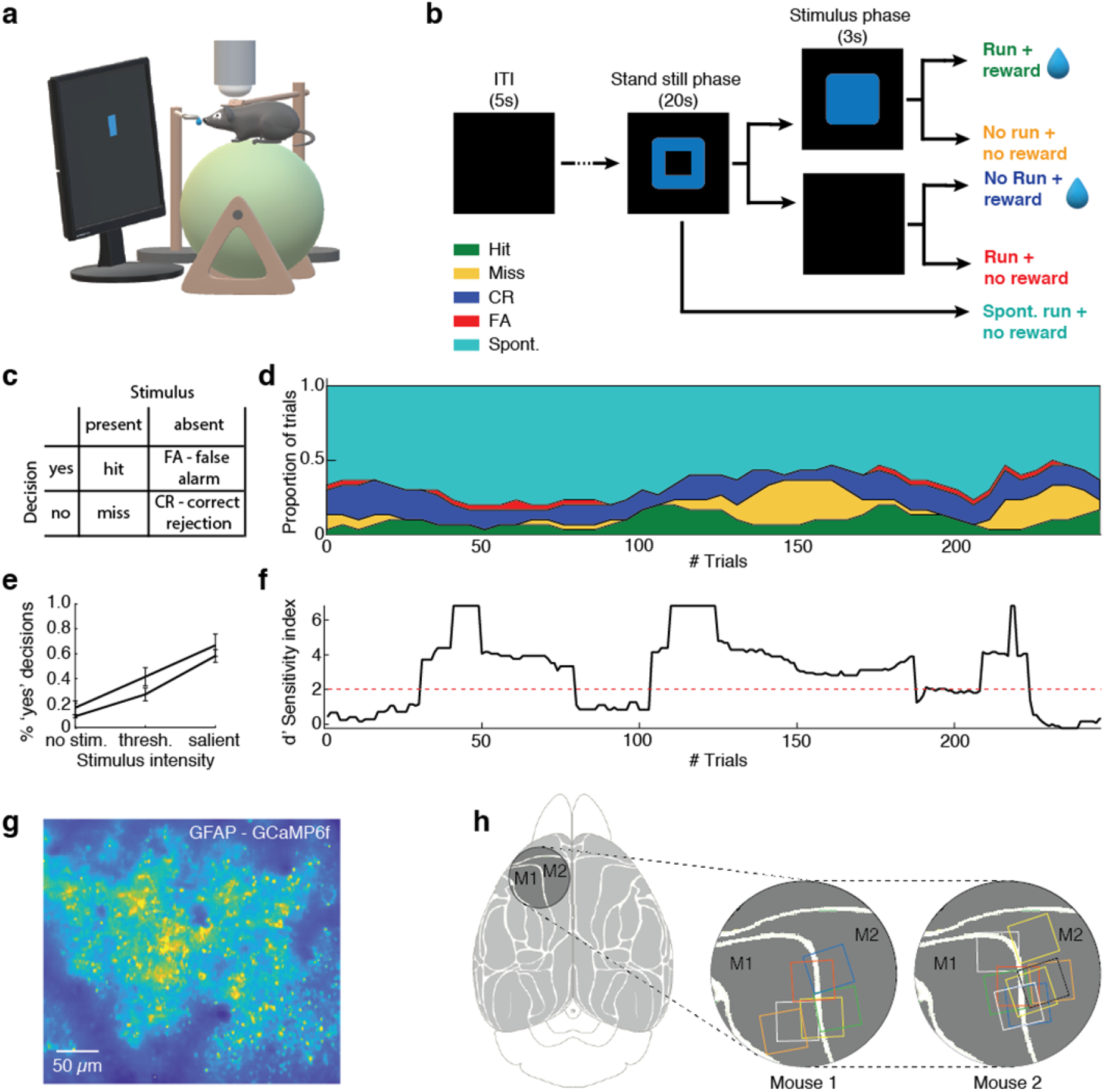
Approach for relating astrocyte syncytium calcium signals to behavioral context using detection task variables. **a**, Schematic of the experimental setup. Head-fixed mice were placed on a spherical treadmill viewing a computer screen. Astrocytic calcium excitation was recorded in layer 2/3 of the M1/M2 motor cortex using two-photon microscopy while the mice performed the visual detection task. In total, we recorded 4,837 trials during 21 behavioral sessions (see **Methods**). **b**, Schematic of the behavioral protocol. A trial started when mice stopped running for 1 s. A visual cue (blue frame) instructed mice to remain still for 20 s. Following this stand-still phase, a stimulus (blue square) was presented for 3 s in 50% of the trials. In the other half of the trials, no stimulus was shown. Stimulus intensity varied between two levels: salient and close to the perceptual threshold (see **Methods**). Stimulus presence and intensity were randomly selected. In trials with stimulus presentation, mice were required to start running within the 3 s stimulus phase to receive a fluid reward. In trials without stimulus presentation, mice had to remain still for a 3 s period to receive the reward. Spontaneous runs during the 20 s stand-still phase aborted stimulus presentation. Mice were able to initiate a new trial after a 5 s inter-trial interval. **c**, Signal detection theory classes for behavioral outcomes (hit, miss, correct rejection, and false alarms), given two stimulus conditions (stimulus present or absent) and two possible decisions (‘yes, stimulus present’ and ‘no, stimulus absent’). **d**, Proportions of behavioral outcomes during one example session. **e**, Average psychometric detection curves for two representative mice. **f**, The mouse’s performance levels during the example session shown in *d*. The performance level was quantified using the d′-sensitivity index, calculated as the difference of z- scores for ‘hit’ and ‘false alarm’ rates. A d′-value of 2 was chosen to distinguish between high- and low-performance states. **g**, Heatmap of average GCaMP6f fluorescence in layer 2/3 from an example recording in area M1. **h**, *Left*, dorsal view of the mouse cortex with the chronic cranial window location indicated (circle). *Center* and *right*, imaging locations (squares) within the cranial window for two representative mice. M1, primary motor cortex; M2, secondary motor cortex.

Astrocyte calcium activity in fully trained GFAP-GCaMP6f mice was recorded using two-photon imaging (**Fig. 1g**). All recordings were performed in cortical layer 2/3 of the primary and secondary motor areas (M1/M2) and had a ∼510×640 µm field-of-view recorded at ∼30.9 Hz (**Fig. 1g,h**). While the GFAP promoter drives expression in most and predominantly astrocytes, a limited region-dependent neuronal expression has been reported^14, 15^. We, therefore, computationally identified and excluded any areas showing features of neuronal activity^16^ (see **Methods**).

Next, we analyzed the animals’ task-related running responses. During hit trials and false alarms (FA), the runs started shortly after stimulus presentation (mean reaction time of two representative mice: 0.7 s and 1.2 s) (**Fig. 2a**, top panel). During correct rejections (CR) and miss trials, mice remained still on the treadmill during the stimulus phase. However, during reward consumption, 97.6% of CR trials were followed by a run. Similarly, stimulus offset triggered runs in 7.2% of the miss trials.

**Fig. 2.**
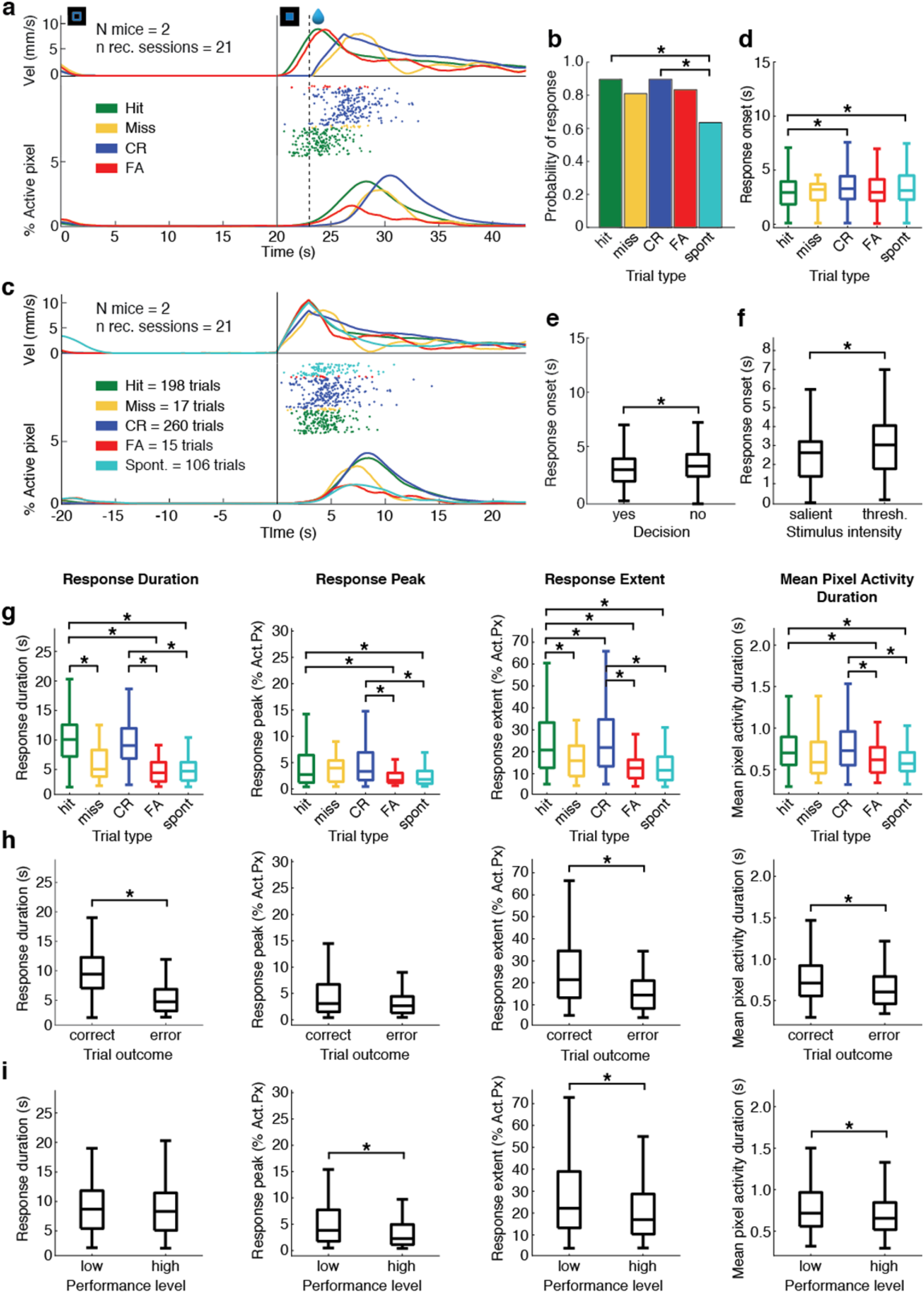
Astrocyte syncytium responses encode detection task variables. **a**-**i,** Astrocyte syncytium responses encode reward, the animal’s decision (stimulus present or stimulus absent), and performance level. **a**, Population data showing the astrocyte syncytium signals’ dependence on the trial type. *Top*, running velocity profile for hit (green), miss (yellow), CR (blue), and FA trials (red). *Center*, onsets (colored dots) for individual qualifying astrocyte syncytium signals by trial type. *Bottom*, average astrocyte syncytium calcium signals, represented as the percentage of ROA (Regions of Activity) pixels over time (see **Extended Data Fig. 1**). Each colored trace is an average across the individual trials of a given type aligned to the stand-still cue onset (198 hit, 17 miss, 260 CR, 15 FA trials, and 106 spontaneous runs from 21 recording sessions). Only trials that included a run within a defined parameter range were included to ensure comparability (see **Methods**). Vertical lines at 20 s and 23 s mark the stimulus phase. **b**, Probability of observing a significant astrocyte syncytium response for the different detection task trial outcomes and spontaneous runs. Only spontaneous runs that occurred 15 s after stimulus onset and before the end of the 20 s stand-still phase were included in the analysis. **c**, Same population data as in *a*, but aligned at run onset (0 s). **d**, Astrocyte syncytium response onsets relative to run onset for the different trial types. The boxplot marks the median and the 25th and 75th percentiles of the data for each trial type. The whiskers cover ∼99.3% of the data. **e**, The animals’ ‘yes’ decision (based on hit and FA trials) was encoded by an earlier onset of the astrocyte syncytium response. **f**, Stimulus intensity was encoded by astrocytes’ syncytium response onsets. **g**, Astrocyte signal strength, as quantified by response duration, peak, the total area under the response curve, and mean pixel activity duration (from left to right), was significantly larger for rewarded than spontaneous and in some characteristics for error trials. **h**, Encoding of rewarded versus error trials. Same layout as in *g*. Rewarded trials showed significantly longer response durations. **i**, The animal’s performance level was encoded primarily by the astrocyte syncytium response amplitude. Low-performance periods were associated with higher amplitudes. The layout is the same as in *g*. Statistical significance was derived from linear mixed-effects model (LME) analysis for all comparisons (see **Methods, Tables 2-10**).

To study astrocytes’ encoding of complex cognitive functions in the context of running, we only analyzed trials that included runs with comparable characteristics (for trial selection criteria and numbers, see **Methods** and **Table 1**). Moreover, we applied multivariate analysis to explore the joint influence of cognitive and run-related variables and determine the effect of each variable in the presence of the others. To analyze astrocytes’ response properties, we implemented a Region of Activity (ROA) analysis algorithm that uses three-dimensional filtering and noise-based thresholding on individual pixels over time to detect significant fluorescence transients^7^ (see **Methods**). Astrocytic syncytium responses were plotted as the percentage of active pixels within the labeled area over time (**Extended Data Fig. 1a**). We characterized the syncytium responses by calculating their onset (relative to run onset), duration, peak value and time, and offset (**Extended Data Fig. 1b**). We also calculated the total extent of activation (i.e., projection of active pixels throughout the response interval normalized to the total labeled area) and mean duration of pixels activated during the response interval. To identify the contribution of behavioral variables (trial type, performance level, recording area, mouse identity, current and previous run parameters) to astrocyte syncytium response characteristics, we used multivariate linear mixed-effects (LME) models with recording sessions as a random effect (see **Methods**).

We found that behavioral context significantly influenced the astrocyte syncytium response to running. Not every run was capable of triggering an astrocyte response. In areas M1/M2, we found a significant effect of inter-run interval period on the probability of eliciting an astrocyte response (**Extended Data Fig. 2a**), a cell-intrinsic mechanism previously reported for cerebellar Bergmann glia^10^. The shorter the rest period, the less probable (**Extended Data Fig. 2b**), weaker (**Extended Data Fig. 2d**), and more delayed astrocyte responses were (**Extended Data Fig. 2c**). Nevertheless, the trial type had a significant effect on the probability of astrocyte syncytium responses. To further investigate this trial type effect, we focused on task-related runs with ≥20 s stand-still phase (by task design) and spontaneous runs with ≥15 s inter-run distance. Notably, astrocyte syncytium response probability was significantly higher for rewarded runs than spontaneously initiated runs (**Fig. 2b**).

Next, we aligned all run trial transients at the run onset to examine how syncytium responses depended on behavioral context parameters. This representation also allowed us to compare task-related trials to spontaneous runs (**Fig. 2c**). Averaged syncytium transients lasted ∼10 s, and their onset latency (3 s) was strongly correlated to run onset (**Fig. 2d**). We found significantly shorter onset latencies (2.7 s) for hit compared to CR trials (3.2 s) and spontaneous runs (3.2 s) (**Table 3**). Additionally, our LME analysis revealed that astrocyte syncytium signals significantly encoded the detection decision, with earlier onsets for hit and FA trials (’yes’ decision) (**Fig. 2e**, **Table 9**). Applying the LME model on hit trials only, we found that syncytium responses to salient stimuli were shorter (2.6 s) than to threshold stimuli (3 s) (**Fig. 2f**,

**Table 10**). The strength of astrocyte syncytium responses (i.e., its response duration, peak, total extent of activation, and mean pixel activation duration) was similar for rewarded trials (hits and CRs) (**Fig. 2g**) but stronger compared to spontaneous runs. Astrocyte syncytium responses also significantly differed between rewarded and error trials, with correct trials showing longer response durations, larger total extent, and longer mean pixel activation duration (**Fig. 2h**).

Astrocyte calcium activity also significantly encoded the animals’ performance levels throughout the session. The response peaks, total extent, and the mean pixel activation duration were significantly larger/longer during low-performance periods than high-performance phases (**Fig. 2i**). Finally, responses in area M1 showed more prominent peaks, total extent, and mean pixel activation duration than in area M2 (**Extended Data Fig. 3**). In summary, astrocyte syncytium responses are extraordinarily versatile, with different response characteristics encoding various behavioral features.

**Fig. 3.**
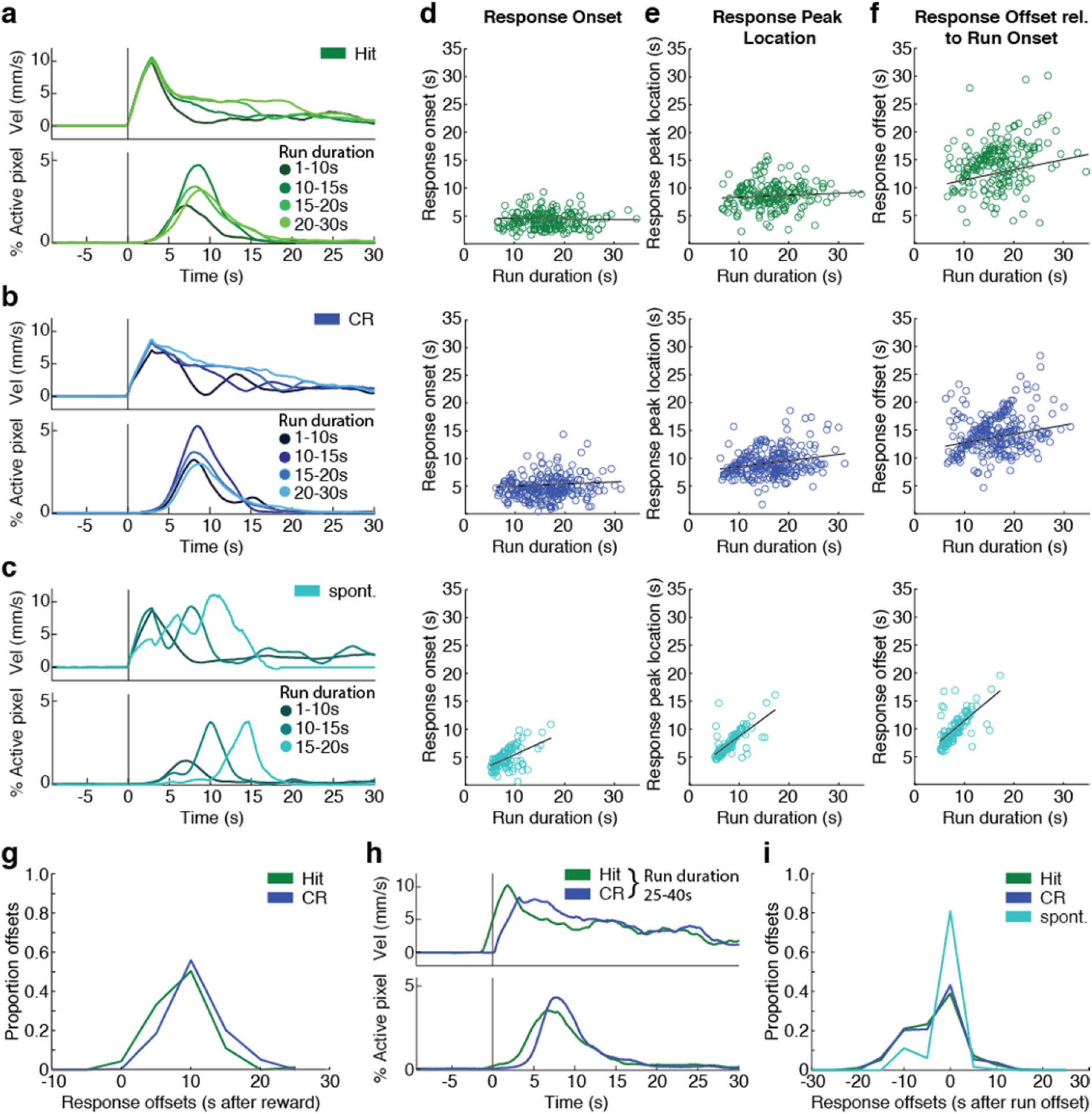
Astrocyte syncytium responses show different run encoding mechanisms. **a**-**c**, Astrocyte syncytium responses, grouped by different run durations, revealed different response profiles for rewarded run trials than spontaneous runs. **a**, Response profile for hit trials. *Top*, running velocity profiles. *Bottom*, astrocyte syncytium responses for hit trials of different run duration (19, 56, 75, and 47 trials of 1-10 s, 10-15 s, 15-20 s, and 20-30 s run duration, respectively, from 21 recording sessions). **b**, Response profile for CR trials. Same layout as in *a*. The data is an average across 36, 73, 98, and 53 runs of 1-10 s, 10-15 s, 15-20 s, and 20-30 s duration, respectively, from 21 recording sessions. **c**, Response profile for spontaneous runs. Same layout as in *a*. The data is an average across 104, 19, and 3 runs of 1- 10 s, 10-15 s, and 15-20 s duration, respectively, from 21 recording sessions. **d**, Astrocyte syncytium response onsets as a function of run duration for hit trials (*top*), CR trials (*center*), and spontaneous runs (*bottom*). **e**, Peak location of the astrocyte syncytium response as a function of run duration. Same layout as in *d*. **f**, Astrocyte syncytium response offsets relative to run onsets as a function of run duration. The layout is the same as in *d*. **g**, Histogram of astrocyte syncytium response offsets for rewarded trials (hit and CR) relative to reward onset (see also **Extended Data Fig. 4d**). Event frequencies were bin-normalized for each run duration interval. **h**, Profile of astrocyte syncytium responses for the longest runs (25-40 s). *Top*, running velocity. *Bottom*, astrocyte syncytium responses aligned at reward onset. **i**, Histogram of astrocyte response offsets relative to run offsets for hit trials, CR trials, and spontaneous runs. Event frequencies were bin-normalized for each run duration interval.

Notably, our LME model also revealed a significant dependence of the astrocyte syncytium response duration on the current run duration with longer runs resulting in slightly longer response durations (19%, 12%, and 35% slope for hits, CRs, and spontaneous runs, respectively; **Extended Data Fig. 4a**, **Table 12**). To examine this dependency more closely, we plotted astrocyte syncytium responses for different run durations (1-10 s, 10-15 s, 15-20 s, 20- 30 s) for the hit and CR trials and spontaneous runs (**Fig. 3a-c**). For rewarded trials, run duration did not affect response onset. In contrast, longer run durations for spontaneous runs resulted in longer response latencies (42% slope, **Fig. 3d**, **Table 13**). Likewise, while the response peak location shifted only slightly toward later time points for longer runs in hit (4% slope) and CR trials (11% slope), we found a considerable peak location shift (66% slope) for spontaneous runs (**Fig. 3e**, **Table 14**). Additionally, when we calculated response offsets relative to the run onset, this duration increased only slightly with run duration for rewarded trials (18% increase for hit trials and 16% for CR trials). In comparison, it changed drastically for spontaneous runs (75% slope) (**Fig. 3f**, **Table 15**). These findings imply that different mechanisms control the on- and offset of astrocyte syncytium responses in different behavioral contexts.

**Fig. 4.**
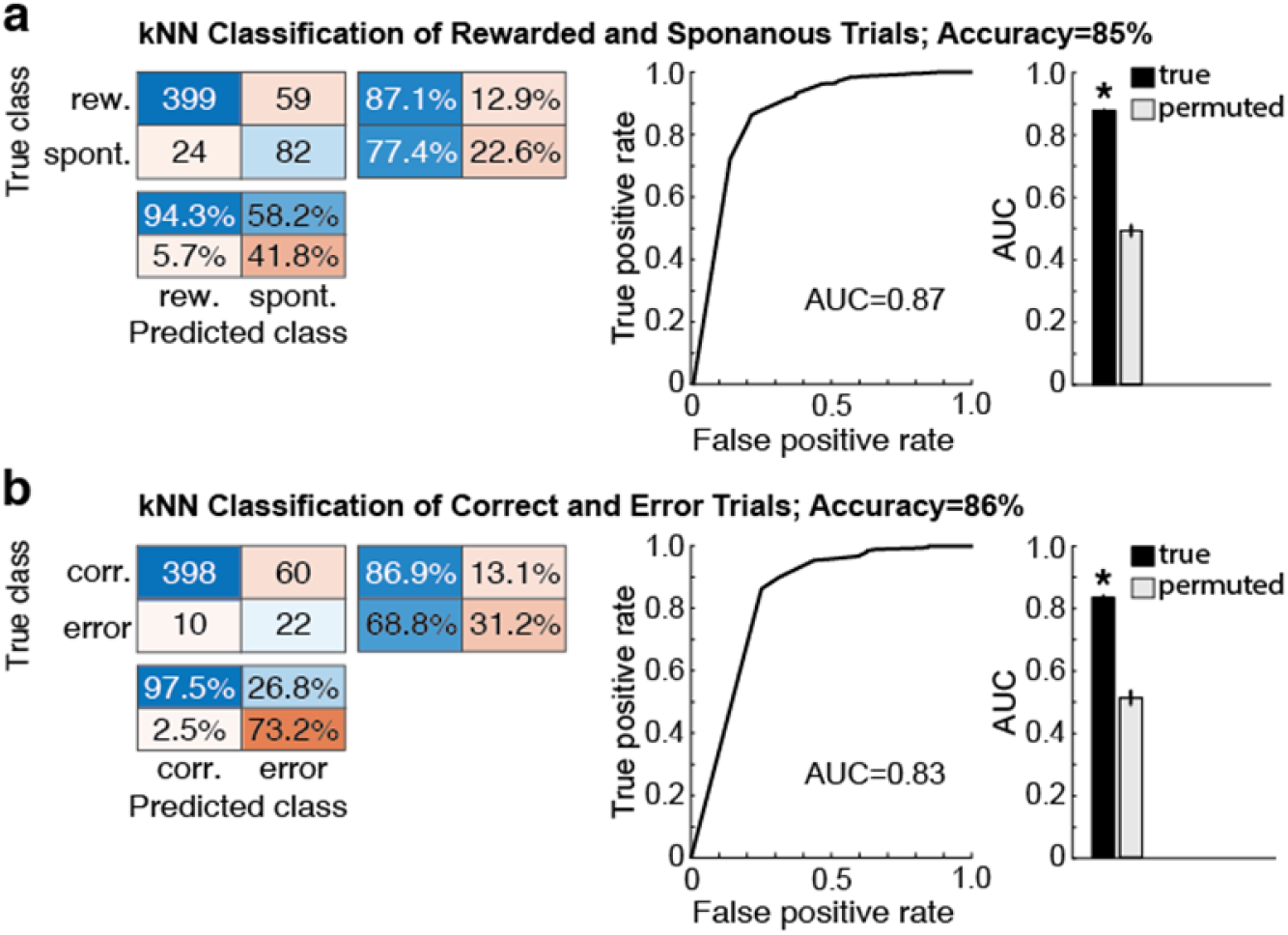
Detection task variables can be decoded from astrocytic syncytium responses using machine learning approaches. **a**-**b**, The k-nearest neighbor (kNN) classifier allows reliable decoding of rewarded/correct trials from astrocytes’ syncytium calcium responses. **a**, Classifier decoding performance of rewarded trials from rewarded trial and spontaneous run astrocyte syncytium responses. *Left*, classifier confusion matrices with rows representing the true classes and columns showing the classifier predictions. The main diagonal shows how frequently the classifier correctly assigned the trials to their real category (accuracy). Off-diagonal cells correspond to the count of incorrectly classified trials. A row-normalized row summary and a column-normalized column summary display the percentages of correctly and incorrectly classified trials for each true class or predicted class, respectively. *Center*, receiver-operating characteristic (ROC) curve and area under the ROC curve (AUC) for the classifier’s output. *Right*, true data mean AUC values (black) were obtained using a 10-fold cross-validation design, repeated 100 times, and compared to the mean AUC values from shuffled trials (gray) when syncytium responses were randomly assigned to one of the two classes. **b**, Classifier decoding performance of rewarded trials from rewarded and erroneous trial syncytium responses. The layout is the same as in *a*. Error bars indicate s.e.m.

Rewarded runs appeared to have a defined onset and offset period of ∼15 s, within which the response peak and duration varied slightly. We calculated the difference between the response offset and run offset to investigate whether a behavioral event might trigger the astrocytic syncytium response offset. For rewarded trials, the response offset coincided with run offset for 13-15 s-long runs (**Extended Data Fig. 4b**, top and center panels). For shorter runs (<13 s), the response duration outlasted the run, and for longer runs (>15 s), it was shorter (**Table 16**).

Intriguingly, this 13-15 s response interval corresponds well with the duration that dopamine is detectable in the extracellular space during rewarded trials^17^ (**Extended Data Fig. 4d**). Both rewarded trial types showed higher response peaks at this ‘preferred’ run duration (**Fig. 3a,b** and **Extended Data Fig. 4c**, top, center panels, **Table 17**). Calculating the response offset relative to the reward onset in correct trials showed that most responses ended ∼10 s after reward onset, with only a few lasting longer than 15 s (**Fig. 3g**). Similarly, aligning the responses to the longest runs (25-40 s) at reward onset showed that their offsets are similar (**Fig. 3h**). In contrast, for spontaneous runs, the astrocyte response co-varied with run duration (**Fig. 3c**), with the difference between syncytium response offset and run offset clustering around 0 s, irrespective of run duration (**Extended Data Fig. 4b**, bottom panel, **Table 16**). As expected, a histogram of response offsets relative to run offset reflects this high degree of correlation (**Fig. 3i**). Together, this data suggests that different encoding profiles underlie rewarded and spontaneous runs and that reward-related molecular signals, such as dopamine, modulate astrocytes’ run-evoked syncytium responses.

While linear regression analysis is restricted to predefined signal characteristics (e.g., onset, offset, peak, or duration), decoding models can access all information contained within the signals’ time course. To infer relevant parameters from the signals’ temporal dynamics, we used the k-nearest neighbor (kNN) classifier^18^, one of the most popular supervised machine learning algorithms for time series classification. We trained the classifier on example syncytium traces, represented as vectors in multidimensional feature space with corresponding class labels. In the subsequent test phase, the classifier was tasked with predicting the classes of syncytium transients that the classifier had not used for learning, based on the most frequent class among the *k* training samples nearest to the query vector. Bayesian optimization was used to select the distance calculation method and *k*, the number of neighbors (**Table 18**). We visualized classifier predictions using confusion matrices^19^. To evaluate the classifier’s performance, we calculated the area under the receiver operating characteristic curve (AUC). This curve captures the true positive versus the false positive rate of the classifier at different classification thresholds, thereby representing the prediction performance quality irrespective of the chosen threshold. Statistical significance was derived from random permutation testing, shuffling the training data class labels, and calculating the probability that the prediction performance could be explained by chance (**Table 19**).

When we used syncytium responses for rewarded and spontaneous trials only, the kNN classifier was able to identify these two classes with high accuracy (85% correct class assignments; chance level at 50%) and AUC=0.87, significantly different from the mean calculated in the permutation test (AUC=0.5, **Fig. 4a**). The classifier also confirmed the high predictability of correct and error trials from syncytium responses (86% accuracy; chance level at 50%; AUC=0.83, **Fig. 4b**). Remarkably, the syncytium response carried information about every trial type, which the classifier could predict from the recorded trials (38% accuracy; chance level at 20%) (**Extended Data Fig. 5a**). Moreover, the classifier predicted the animals’ performance level using all traces from all recorded trial types (62% accuracy; chance level at 50%; AUC=0.64, **Extended Data Fig. 5b**).

Next, motivated by our LME model results showing that astrocyte syncytium responses varied substantially with run duration for spontaneous but not as much for task-related runs, we asked whether the kNN classifier could predict run duration from spontaneous runs (**Extended Data Fig. 5c**) and task trials (**Extended Data Fig. 5d**). We found that decoding of run duration was possible from both spontaneous (90% accuracy; chance level at 33%) and task trials (46% accuracy; chance level at 25%), with significantly higher accuracy and AUC values when spontaneous trials were used for classification (p<0.05, Kolmogorov-Smirnov test). We reasoned that if the gradual increase of run duration in the defined run duration classes is accompanied by a gradual change in the encoding signal, the classifier decoding performance should be most robust along the main diagonal of the confusion matrix, and confusions between adjacent classes should be more frequent. To test this idea, we averaged the classification probability along the main diagonal and the parallel diagonals, resulting in the average performance of the classifier as a function of distance from the actual run duration (**Extended Data Fig. 5c-d**, last panel). The function peaked at the probability for correctly assigning the query traces to their real run duration class, while more erroneous classifications occurred for adjacent run durations. This finding confirms the gradually changing nature of the signal underlying the decoding of spontaneous and task trials and the proper operation of the classifier.

While the previous analyses demonstrated that we could decode animal behavior (run context, reward delivery, performance level, run duration) from astrocyte syncytium responses, we also wanted to know whether the responses were relevant for the animals’ behavior. If the astrocytic signal is relevant for mouse behavior, the decision should be decodable from correct and error trials. Indeed, we found that the perceptual decision of the animal could be decoded from correct (hit and CR) trials (62% accuracy; chance level at 50%; **Extended Data Fig. 6a**). Next, we trained the kNN classifier on correct decision trials (hit trial: ’yes’ decision; CR trial: ’no’ decision) and used the signals for erroneous decisions (miss trial: ‘no’ decision, FA trial: ’yes’ decision) as a test dataset. We found that error trials also carried significant information about the decision (**Extended Data Fig. 6b**). Finally, we examined whether areas M1/M2 encoded information about the nature of the sensory information that was essential for the animals’ decision. In accordance with our LME model results (**Fig. 2f**), the classifier was able to decode information about the presented stimulus intensity from the astrocyte syncytium responses to hit trials (65% accuracy; chance level at 50%; **Extended Data Fig. 6c**). Importantly, decoding of stimulus type information was not possible from astrocyte syncytium responses to miss trials (53% accuracy; **Extended Data Fig. 6d**), implying that sensory information important for decision-making was absent in error trials. Together, these findings suggest that the information encoded by astrocyte syncytium responses is relevant for animal behavior.

## Discussion

In summary, the astrocyte syncytium calcium response is a complex yet deterministic signal encoding several aspects of behavioral context. Signal onset was tightly linked to run onset in rewarded trials, with an earlier calcium response encoding the animal’s decision (**Fig. 2e**) and stimulus intensity (**Fig. 2f**). Interestingly, in spontaneous runs, the response onset had a significant delay for longer run durations. Response duration was influenced by both decision correctness in task trials and run duration (**Fig. 2g**). Response offset correlated with dopamine levels in rewarded trials and run offset in spontaneous runs (**Extended Data Fig. 4d**, **Fig. 3i**). The overall strength of the calcium response was impacted by trial type, with rewarded trials showing the most notable increase (**Fig. 2g**). The amplitude was also significantly modulated by the animal’s performance level (**Fig. 2i**) and potentially by run parameters linked to reward expectation (**Fig. 3a-b**). The inter-run interval had a significant impact on the probability and strength of the astrocytic response in task trials and spontaneously initiated runs. Notably, the information encoded in the astrocyte syncytium calcium responses was behaviorally relevant (**Extended Data Fig. 6**).

What mechanisms might control astrocyte syncytium responses? Because astrocytes do not exhibit stereotyped calcium waveforms like those evoked by neuronal action potentials, previous work suggested that their transients result from spatial and temporal integration of behavior-related extracellular molecular signals released, for example, by local and projection neurons^1^. The complex yet deterministic nature of astrocyte syncytium responses revealed by our study supports this notion. Response duration and amplitude depended, amongst others, on run duration, suggesting integration of ongoing synaptic activity by astrocytes (**Fig. 3**). Rewarded hit and CR trials showed larger syncytium responses than unrewarded trials with a ’preferred’ run duration. One possible explanation for this ‘preferred’ run duration is that 13 s-long runs offer the highest reward probability to the animal, with astrocytes reflecting the corresponding local activity of M1/M2 neurons. Another possibility is that dopamine’s time course determines the peak of the astrocyte syncytium response in rewarded runs. The time course of previously measured dopamine signals in the same region and task are consistent with this hypothesis^17, 20^ (**Fig. 2**; **Extended Data Fig. 4**). We also found that astrocyte syncytium responses in run trials are significantly different from no-run trials. Specifically, the probability of miss and CR trials without a run was significantly lower than those with a run (**Extended Data Fig. 7a,b**).

Moreover, the syncytium responses to no-run trials had significantly longer response latencies (**Extended Data Fig. 7c**), were shorter, reached lower peak values, and showed lower total activation extent (**Extended Data Fig. 7d**). This observation seems consistent with previous work showing that locomotion mediates noradrenaline release and widespread astrocyte calcium excitation and that the astrocyte response is boosted in the presence of sensorimotor- evoked local neural activity^11, 12^. How astrocyte syncytium responses may differ in behavioral tasks that do not involve a running response (e.g., lever press/release task) remains to be determined. Apart from dopamine and noradrenaline, additional neuromodulator signals, such as acetylcholine, may also modulate astrocytes’ phasic syncytium responses^21, 22^. Our finding that astrocyte responses were larger during low-performance states may, at least in part, be explained by higher tonic neuromodulator levels (e.g., noradrenaline) associated with this cortical state^23, 24^ (**Fig. 2**). Together, our data seem consistent with the concept of spatial and temporal integration of neurotransmitter and neuromodulator signals in shaping astrocyte syncytium responses in the M1/M2 cortex. Nevertheless, to better understand the syncytium signal’s building blocks and regional differences, an analysis of individual regions of interest or astrocyte compartments may be informative.

How might astrocyte syncytium responses affect local neural circuits? Our findings suggest that astrocytes’ signaling is encoding relevant behavioral information (**Extended Data Fig. 6**).

However, one of the striking features of astrocyte syncytium responses is their seconds-long delay relative to run onset (**Fig. 2**; **Extended Data Fig. 1**), likely due to signal integration within astrocytes (e.g., IP3, which mediates endoplasmic reticulum (ER) calcium release) ^3, 25^. This delay, together with the syncytium responses’ slow time course, indicates that astrocyte excitation likely serves complementary roles to neuronal activity, particularly those preceding or initiating the behavioral response (e.g., decision making or motor planning). One potential role of astrocyte syncytium responses may be circuit regulation. Following task execution, astrocytes may restore the neural circuit’s ionic and transmitter homeostasis, thus ensuring the circuit’s continued operation with optimal signal-to-noise ratio and gain. Additionally, they may actively tune the system when the executed behavior does not reliably achieve the desired outcome (e.g., reward)^26, 27^. By establishing a computational “review period” of past events, astrocytes could potentially inform future behavior, enabling trial-to-trial behavioral adjustments or learning. If these considerations are correct, they might explain why astrocyte syncytium responses depended on behavioral performance and perceptual level (**Figs. 1-2**). These hypotheses might be tested by an in-depth analysis of trial history and performance as a function of the inter-run interval, which strongly affects astrocytes’ response probability and strength (**Extended Data Fig. 2**).

Given the complex yet predictable syncytium responses (**Fig. 4**; **Extended Data Figs. 5-6**), it is conceivable that astrocyte calcium excitation mediates more than one output and on multiple timescales. Astrocytes can modulate neural circuit activity on the seconds (i.e., individual trial) timescale by releasing neuroactive substances in a calcium-dependent manner (e.g., ATP/adenosine, D-serine, potassium)^3^. Neural circuit activity can also be modulated on the minutes (i.e., performance level) timescale by activity-dependent changes in astrocyte transporter activity, gap junctional coupling, metabolic support, or perisynaptic process structure.

How might these open questions about astrocyte syncytium responses be addressed? The quantitative visual detection task and computational methods employed in our study may help address these fundamental questions. In conjunction with genetically encoded neurotransmitter and neuromodulator sensors, our standardized approach may help reveal how behavior-dependent extracellular signals relate to astrocyte activity, as exemplified for dopamine (**Extended Data Fig. 4**). However, this may require further optimization of current transmitter sensors and their color variants to enable concurrent and high-resolution measurement of corresponding transient maps. Simultaneous recording of astrocyte and projection neuron activity can only partly replace such measurements, as calcium spiking does not identify the type and quantity of the transmitter(s) released or its spatial spread. Likewise, new indicators for intracellular signaling (e.g., IP3, cAMP, or PKA) and functional alterations (e.g., proximity assays) may in the future allow measurement of how the various molecular signals are integrated within astrocytes, how this spatiotemporal integration relates to astrocyte syncytium responses, and how these responses modulate astrocyte output^28, 29^. One approach to determine the effect of astrocyte syncytium responses on local neural activity may be to leverage their intrinsic properties. We showed that the probability, onset, and magnitude of syncytium responses depend on inter-run distance (**Extended Data Fig. 2**), an effect previously described for cerebellar astrocytes and likely dependent on ER calcium store dynamics^10^.

Animals trained to perform visual detection task trials at various inter-run distances may provide insight into how local neural activity changes in the presence or absence of astrocyte syncytium responses. However, the dependency of astrocyte syncytium responses on trial-type, performance levels, and other behavioral variables suggests that approaches to globally in- or decrease astrocyte excitation (e.g., by opsin, DREADD, calcium pump, or chelator expression) may only partially mimic astrocytes’ varied effects on neural circuits. Finally, applying our visual detection task and computational methods to other (e.g., sensory) brain regions should help determine conserved features of astrocyte encoding and circuit modulation and inform models of how astrocyte signaling may need to be incorporated into systems neuroscience.

## Methods

### Experimental model and subject details

All procedures were performed following the National Institutes of Health (NIH) guidelines for the Care and Use of Laboratory Animals and were approved by the Institutional Animal Care and Use Committee (IACUC) at the Salk Institute. Mouse strains used in this study included Gfap-Cre 73.12 (RRID: IMSR_JAX:012886) and Ai95D mice (RRID: IMSR_JAX:024105)^30, 31^.

All imaging and behavioral experiments involved heterozygous male mice (N=4). Mice underwent two surgeries: head plate implantation at 8-10 weeks of age and cranial window implantation at ∼12 weeks of age. Training started ∼7 days after each surgery. Mice were water-restricted to 25 ml kg^-1^ per day and maintained at 80-85% of their normal ad-libitum weight during training and imaging. Optical recordings were performed at ∼20 weeks of age. Of the five mice trained on the task, one failed to reach proficiency. Mice were typically group-housed, provided with bedding and nesting material, and maintained on a 12 h light-dark cycle in a temperature (around 22**°**C) and humidity controlled (45-65%) environment. The animals had ad libitum access to standard rodent chow and water outside of training and imaging periods.

### Live animal preparation

Head plate and cranial window implantation were performed as previously described^17, 20^. Briefly, mice were anesthetized with isoflurane (4% and 2% for induction and maintenance, respectively) on a custom surgical bed (Thorlabs Inc., Newton, NJ). Body temperature was maintained at 36–37°C with a DC temperature control system. Ophthalmic ointment was used to prevent eyes from drying. The skin at the surgical site was cleaned and disinfected with 70% ethanol and Betadine. A small (∼10 mm) incision was performed along the midline. The scalp was pulled open, and the periosteum was cleaned. A portion of the scalp was surgically removed to expose frontal, parietal, and interparietal skull segments. A custom metal plate was affixed to the bone above the motor cortex with C&B Metabond Quick Adhesive Cement (Parkell Inc., Edgewood, NY). The cement also covered all other exposed skull regions.

After initial training, a custom-made cranial window was implanted to enable chronic two-photon imaging^32^. The skull was thinned above the motor cortex, and a craniotomy was performed (2.5 mm diameter; centered around AP 1.5 mm / ML 1.5 mm). The dura mater was kept intact. The craniotomy was sealed with a custom three-layered cover glass assembly (each No.1 thickness) with the two layers closest to the cortex consisting of two circular 2.5 mm-diameter coverslips and the outermost layer consisting of a circular 3 mm-diameter cover glass that rested on the thinned skull. UV-curing optical adhesive (NOA 71, Norland Products, Inc.; cat. no. 7106) was used to attach the coverslips one at a time, taking care to avoid air inclusions that might interfere with imaging or facilitate cover glass detachment during the implantation period.

### Behavioral setup and data acquisition

Animal training was performed in a sound-attenuating cubicle (ENV-017M, Med Associates Inc.) using a custom-built setup. This setup included a color LCD monitor for stimulus presentation (12.1“ LCD Display Kit/500cd/VGA, ICP Deutschland GmbH). Noise in optical recordings was minimized by covering the monitor with a color filter (R342 Rose Pink, Rosco Laboratories Inc.). The setup also included a spherical treadmill (Habitrail Mini Exercise Ball, Animal World Network), allowing the animal to run freely or when instructed. Mice were placed on the treadmill facing the LCD display. Head fixation was achieved by clamping the head plate with custom-build holders. An optical encoder (E7P OEM, US Digital) attached to the treadmill enabled measurement of both speed and direction of ball movement. Water reward was delivered with a programmable syringe pump (NE-500 OEM Syringe Pump, New Era Pump Systems, Inc.).

Behavior-related signals were acquired through a data acquisition board (PCI-6221, National Instruments) connected to a breakout box (BNC-2110, National Instruments) and interfaced to MATLAB using the Data Acquisition Toolbox (Version R2010bSP2, The MathWorks Inc.). The MATLAB-based software MonkeyLogic (www.monkeylogic.net)^33,34^ controlled the behavioral task sequence. Custom-written functions were added to MonkeyLogic to enable analysis and control of ball rotation parameters. Treadmill encoder signals and trial marker codes, generated by MonkeyLogic, were acquired (10 kHz sampling rate; ±5 V input range) in sync with the imaging data. Simultaneous acquisition through the microscope’s software (MScan; Sutter Instrument Company) allowed run parameters, behavioral task events, and image frames to be linked with high temporal precision.

### Animal training

Mice were handled/tamed on two consecutive days before behavioral training to reduce stress. During the first two training days, mice spent ∼15-30 min/day in the setup to become accustomed to head restraint. Mice were then trained daily for 60-90 min during which they performed ∼300-700 trials. A sequence of trial task events was initiated when mice stood still on the ball for 1 s. First, a blue square frame was displayed on the monitor, requiring the animal to continue standing still for 20 s (ball rotational velocity ≤2 mm/s). If the mouse remained still for this entire stand-still phase, a second stimulus (filled blue square) was presented for 3 s in 50% of trials, instructing the mouse to initiate a run. The stimulus was presented at two intensities: salient or close to the perceptual threshold (determined empirically towards the end of the training and kept at the same level during recordings). If the mouse initiated sustained movement during the 3 s stimulus phase (ball rotational velocity >10 mm/s for at least 1 s), a water reward was delivered (hit trial). If no running occurred, the trial counted as a miss trial. In the 50% of trials where no stimulus was presented, the mouse received a fluid reward when it remained still on the ball for 3 s (correct rejection, CR). Running during the 3 s period was counted as a false alarm (FA) trial. The trial was aborted and counted as a spontaneous run if the animal moved during the 20 s stand-still phase (ball rotational velocity >2 mm/s). The mouse could initiate a new trial after an inter-trial interval (ITI) of 5 s.

### In vivo two-photon imaging

Once mice had reached proficiency on the task, two-photon imaging commenced. Mice were imaged daily for 9-12 days while performing the task. We used a resonant scanning two-photon microscope (Sutter Instrument) equipped with a pulsed femtosecond Ti:Sapphire laser (Chameleon Ultra II, Coherent) for simultaneous optical and analog data acquisition. GCaMP6f fluorescence was excited with 910 nm light and detected using an ET525/70M emission filter (Chroma Technology Corp.) and H7422-40 GaAsP photomultiplier tube (Hamamatsu Photonics). Average excitation power depended on imaging depth (typically 55-66 mW). The typical recording depth was 100-135 μm below the pia. Data were acquired using a Nikon 16×0.8-NA water immersion objective. A custom-made blackout curtain around the microscope’s detector was used to reduce light contamination by the LCD monitor. Images (512×512 pixels) were acquired at 1.0x Zoom (∼510×640 μm effective field of view after cropping) and ∼30.9 frames/s using MScan software (Sutter Instrument Company). Each recording session consisted of five to twelve ∼10 min recordings, separated by short imaging breaks (3-5 min). Recordings within a given session were performed at the same location to maximize the number of trial repetitions for analysis. Recordings from different sessions on consecutive days were offset either laterally or axially to maximize the tissue volume being sampled (**Fig. 1h**).

### Behavioral data processing and analysis

All data analyses were performed using custom-written MATLAB scripts (The MathWorks Inc). The encoder signal (frequency of voltage changes) was converted to run velocity and smoothed with a 3 s moving average window. This smoothing widened the signal by half the window size. We, therefore, shifted the smoothed velocity trace by +1.5 s for all analyses involving alignments at task events (trial onset, reward onset). Smoothing also lowered peak running speeds (**Fig. 2a**). Run onset threshold was set at 0.5 mm/s, while run offset was defined as the time when running speed fell below 0.1 mm/s. We chose these low thresholds because even small movements could elicit calcium responses. Run onsets <0.5 s after the offset of a previous run were considered as one running event. A trial-associated run counted as hit or FA if running lasted >1 s and exceeded 30 mm/s. The absence of ball movement (<0.5 mm/s) during the stimulus phase counted as CR trial. Run velocity was measured in real-time during animal behavior (without temporal smoothing). While all hit and FA trials included a run during the stimulus phase, miss and CR trials did not exhibit a run during this phase. However, they were often followed by a run after stimulus offset or reward delivery, respectively.

To quantify the animals’ task performance, we recorded all trial outcomes (**Fig. 1c**) and reaction times (RTs) (the time interval between stimulus and run onset). Mice were considered to have reached task proficiency when the proportion of correct decisions (hit and CR trials) exceeded 50% over a 50 trial performance interval. Additionally, RTs for correct ‘yes’ decisions (hit trials) had to drop below 1.5 s. The psychometric curve for each mouse (**Fig. 1e**) was computed based on the proportion of ‘yes’ decisions for stimulus-absent (FA), threshold, and salient stimulus intensity trials (hit trials). A steep increase in miss trials at the end of the session indicated that mice had lost interest in the water reward. Trials beyond that point were excluded from the average performance analysis. To quantify the level of performance throughout the session, we calculated the discriminability index d-prime (d’, **Fig. 1f**) for each session as Z(hit/(hit + miss)) − Z(FA/(FA + CR)), with Z(p), p ∈ [0,1]^13^. All trials during d’>2 phases were considered high-level performance trials. Trials during d’<2 phases were deemed to be low-level performance trials.

### Image data processing and analysis

Lateral image motion (e.g., due to mouse movement) was corrected using the non-rigid movement correction algorithm NoRMCorre^35^. We used 200 frames from the first recording of a given session to compute the registration template. The same reference image was used to correct the image motion of other recordings taken at the same location.

We excluded regions over and immediately surrounding blood vessels to reduce artifacts caused by vascular dilation and constriction. First, we calculated a baseline image by smoothing the image data temporally with a moving average of 1 s. Next, we determined the mode of the pixels. Areas below the 70^th^ percentile of the baseline image’s pixel intensity distribution were automatically excluded from data analysis.

While the GFAP promoter drives expression in most and predominantly astrocytes, a limited region-dependent neuronal expression (0.5-5% of labeled cells) has been described^14, 36^. To identify corresponding regions in our data, we first calculated the mean intensity projection of all recordings at a given imaging site and segmented this image using the CellProfiler image- analysis software^37^. We allowed the total area of segments to vary between 8 and 300 pixels.

Next, we extracted the fluorescence time trace F(t) from all segments by averaging the pixel intensities of all pixels within individual segments. ΔF(t)/F was calculated as (F(t) – mean F) / mean F. Segments were classified manually by considering their morphology (from the mean intensity projection image), waveform shape, and event frequency and pattern (from the corresponding ΔF(t)/F trace). Segments showing features of neuronal activity^16^ were excluded. Between 4.5% and 12.8% of segments displayed neuronal characteristics in areas M1/M2.

To capture the high spatiotemporal complexity of astrocytes’ calcium signals, we implemented a previously described activity-based algorithm based on Regions-of-Activity (ROA) analysis^7^ with a few modifications. The data were smoothed with a Gaussian filter (*σ* = 3 pixels). To remove slow drifts in the calcium baseline, we detrended the time course of each pixel using the MATLAB function detrend() instead of bandpass filtering the data. Fluorescence events were determined based on noise-based thresholding over time for each pixel. First, the signals were high pass filtered. Then, the standard deviation of each pixel’s noise over time was calculated. Whenever a given pixel’s value in the standard deviation image exceeded the corresponding value 5-fold, the pixel was considered active. The syncytium response signal was calculated as the sum of the active pixels in the field of view (FOV) normalized by the total GCaMP6f labeled area over time.

Each astrocyte syncytium time trace includes multiple repetitions of the same trial type. To characterize the syncytium response to a given trial, we quantified its temporal features, such as response onset/offset, probability, and strength. To calculate response onset/offset, we first determined the mean syncytium response distribution during the baseline period (7-2 s before stimulus onset). We defined the 95^th^ percentile of this baseline activity distribution as the significant response threshold (**Extended Data Fig. 1b**). Time points at which the signal surpassed or fell below this threshold relative to run onset were defined as response onset and offset, respectively. Response probability was calculated by determining the proportion of trials during which the astrocyte syncytium signal exceeded the threshold value during the 0.5-15 s response interval after run onset (if a run happened) or 0.5-17 s after stimulus onset (if no run was detected during that trial). Response strength was characterized by the (1) response duration (i.e., the interval between response on- and offset), (2) response peak (the maximum value reached during the response duration), (3) total spatial extent of the syncytium response (defined as the percentage of active pixels in the projection image during the response interval), and (4) mean duration of consecutively active pixels during the response interval.

### Statistical analysis

In total, we analyzed 4,837 trials from 21 behavioral sessions. Only trials with >7 mm/s ball rotational velocity (smoothed) and >5 s run duration were included in our analysis to ensure trial comparability. The analysis also included only spontaneous runs starting >15 s after the stand- still cue onset to ensure comparability of task-trial and spontaneous run-evoked syncytium responses. The resulting numbers of qualifying trials are shown in **Table 1**. Qualifying trial traces from all animals, all sessions, and trial types, associated with a run and significant syncytium response, were averaged and aligned at trial onset (**Fig. 2a**) or run onset to calculate population responses (**Fig. 2c**).

To quantify the relationship between astrocyte syncytium responses and behavioral variables, we performed linear mixed-effects analyses in MATLAB^38^ (**Fig. 2**). Separate encoding models were fitted for astrocyte syncytium response probability, onset, duration, peak (log-transformed), total extent (log-transformed), and the mean duration of pixel activation (log-transformed) as dependent variables. Subject (mouse identity), recording area (M1/M2), trial type (hit, miss, CR, FA, spontaneous run), performance level (high/low), current run duration, current run amplitude, preceding run duration, preceding run amplitude, and the interval between current run onset and preceding run offset were included as fixed effects in the model. The recording session was treated as a random effect. To decide which behavioral variables to include in the model, we first performed a univariate analysis. We included the fixed factors separately and added the random effects to the model. If the p-value of a dependent variable’s relationship to the tested fixed effect was <0.1, the factor was considered for inclusion in the final model. Next, a model with all qualifying fixed effects and the random effect was set up for each dependent variable, followed by a backward step-down model selection. With every iteration, we excluded the fixed effect with the highest non-significant p-value until the p-values of all remaining factors were <0.05. This value was chosen as the significance criterion. Visual inspection of residual plots for the final models did not reveal any apparent deviations from homoscedasticity or normality. We fitted a binomial generalized linear model using the MATLAB function fitglme() (**Table 2**) to analyze the relationship between syncytium response probability and behavioral parameters.

We included only trials followed by a run and compared the proportions of trials with significant syncytium response to the proportion of trials without one (i.e., when the syncytium response remained under the threshold value). We also included the interaction between trial type and run duration in the model because this described our data better (p<0.0001, Likelihood ratio test). For all other dependent variables, we fitted ordinary linear mixed-effects models using the MATLAB function fitlme() and included only trials with a run and significant syncytium response (**Tables 3-10**). To analyze differences between trials with or without a run, we used all trials with a significant syncytium response aligned at trial onset (**Table 11**). Mixed-effects model parameters were estimated by the maximum likelihood method. The significance of the regression coefficients was assessed using the t-statistic.

To investigate the relationship between run duration and syncytium response duration (**Extended Data Fig. 4a**), onset, peak location, and offset, aligned on run onset (**Fig. 3d-f**) or run offset (**Extended Data Fig. 4b**), we also applied linear regression analysis. Linear mixed-effects models were fitted separately for the rewarded trials (hits, CR) and spontaneous runs. Model selection criteria and analysis were the same as described above. The percent slope was determined by multiplying the regression coefficient by 100 (**Tables 12-16**).

To examine the relationship between response peak and run duration, we used responses with peak values >3% active pixels (**Extended Data Fig. 4c**). The peak values appeared to reach their maximum for 13 s-long runs. Linear models were fitted to trials with shorter (<13 s) and longer run durations (>13 s) separately for rewarded trials and spontaneous runs (**Table 17**).

To determine what triggers astrocyte syncytium response offset, we plotted the distribution of rewarded trial offsets aligned at reward onset (**Fig. 3g**). For comparison, we also plotted the rewarded and spontaneous run offsets aligned at run offset (**Fig. 3i**). For both histograms, we normalized the distribution of each predefined run duration interval (5-10 s, 10-15 s, 15-20 s, 20- 25 s, 25 s-maximum run duration). For final representation, the normalized distributions were averaged. This approach helped to avoid biasing run durations that appeared more frequently during trials.

To decode information from astrocyte syncytium responses, we applied the k-Nearest Neighbor (kNN) classifier using the MATLAB function fitcknn() (**Fig. 4** and **Extended Data Figs. 5-6**). We represented the syncytium responses (% active pixels over time), from run onset to 30 s after run onset, as vectors in multidimensional feature space. The prior probabilities for all classes were defined as equal (i.e., 1/number of classes). The classifications were performed in a 10- fold cross-validation design (i.e., data was partitioned in 10 randomly chosen subsets). One data subset was used to validate the model, while the remaining subsets were used for training. We used automatic hyperparameter optimization to find hyperparameters that minimized the 10- fold cross-validation loss (see **Table 18** for resulting parameters for each classification).

Accuracy was calculated as the number of correct predictions divided by the total number of predictions. For decoding the animal’s decision from erroneous syncytium responses, the classifier was trained on correct responses only. It was then tested on error trials that the classifier had not seen before.

To visualize the classifier performance, we used the “Receiver Operator Characteristic” (ROC) curve. We calculated the “Area Under the Curve” (AUC) to measure the classifier’s ability to distinguish between classes. We extended the ROC-AUC calculation to multiclass classification decoding, using the “one versus all technique” (i.e., the ROC for one class was generated to classify this class against everything else).

The classification process and the calculation of the AUC were repeated 100-times to ensure a reliable estimate of the average classification performance. To evaluate the significance of the classifier performance, we used permutation testing. In this test, the response traces were kept the same, but their labels were randomly permutated. After repeating the permutation procedure 100-times, we calculated the AUCs for a classifier trained on a dataset with randomly assigned labels and tested on true classes. This approach generated a null distribution, which we used for the empirical p-value calculation (i.e., the proportion of permutations for which the AUC is greater than the score obtained using the original data)^39^ (**Table 19**).

## Reporting summary

Further information on research design is available in the Research Reporting Summary linked to this paper.

## Data availability

The data that support the findings of this study will be deposited in the Brain Image Library (BIL; https://www.brainimagelibrary.org/index.html), as required for this BRAIN Initiative-funded project. They will also be available from the corresponding authors upon reasonable request.

## Code availability

The custom Matlab-based code used for acquisition, processing, and analysis of the data will be deposited in GitHub, as required for this BRAIN Initiative-funded project. It will also be available from the corresponding authors upon reasonable request.

## Acknowledgments

We thank J.H. Reynolds, T.J. Sejnowski, the U19 A-Team, and members of the Nimmerjahn lab for comments on the manuscript, the Salk machine shop for technical support, and J. Chambers for mouse colony management. This work was primarily supported by the National Institutes of Health (NIH) grant U19NS123719 (A.N.). It was partially supported by the NIH grants U01NS103522 (A.N.), R01NS108034 (A.N) and UL1TR001442 (Altman Clinical & Translational Research Institute; ACTRI). K.M. was a DFG research fellow and Catharina Foundation postdoctoral scholar. The content is solely the authors’ responsibility and does not necessarily represent the official views of the NIH.

## Author contributions

K.M. and A.N. conceived and designed the study. K.M. designed and performed the behavioral and imaging experiments with help from R.W.F. and D.D. K.M. designed and performed the statistical analysis with help from ACTRI services and prepared the figures. K.M. and A.N. wrote the text. All authors discussed the results, provided input or edits on the manuscript.

## Competing interests

The authors declare no competing interests.

Correspondence and requests for materials should be addressed to K.M. and A.N.

## Extended Data Figures

**Extended Data Fig. 1.**
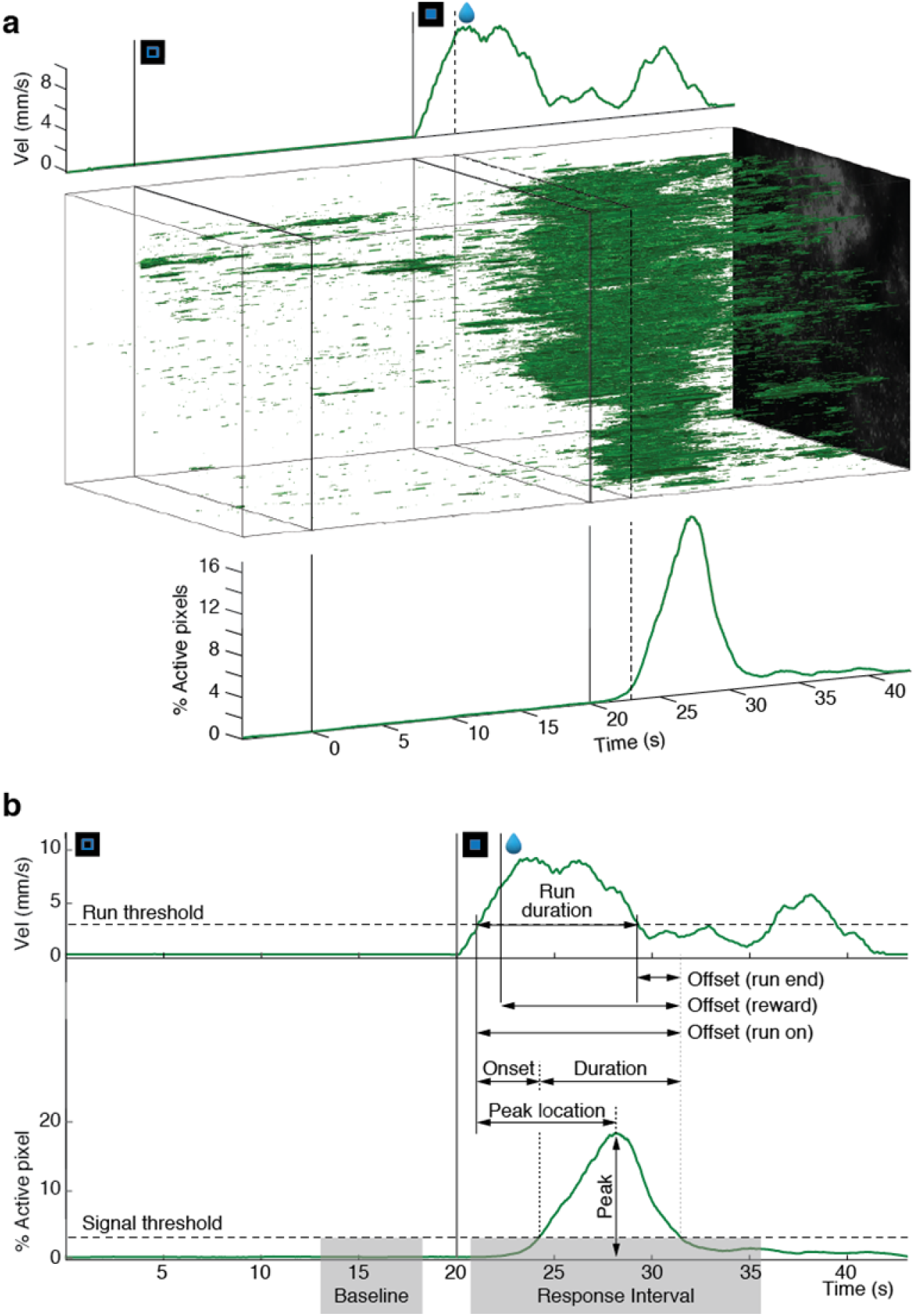
Approach for extracting and analyzing astrocyte syncytium calcium signals. **a**, The Regions of Activity (ROA) algorithm^7^ was used to extract and characterize astrocyte syncytium calcium signals. The example data shows one representative hit trial. *Top*, run velocity profile. *Center*, x-y-t rendering of active pixels detected within the (∼510×640 µm field-of-view (FOV). *Bottom*, the percentage of active pixels over time normalized to all labeled pixels within the FOV. **b**, Schematic of the astrocyte signal characteristics used for data analysis. *Top*, run velocity profile. *Bottom*, astrocyte syncytium calcium signal.

**Extended Data Fig. 2.**
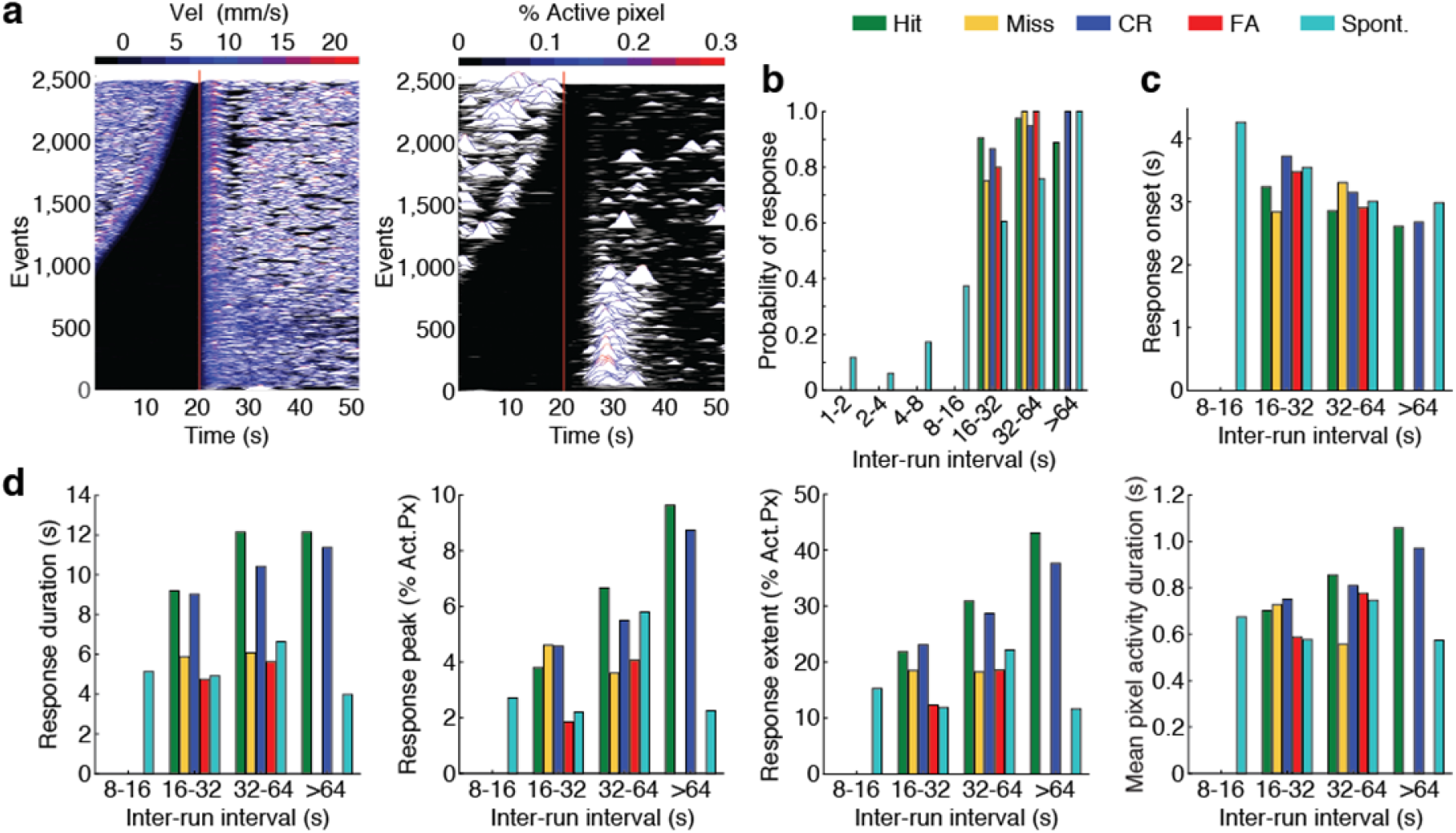
Astrocyte syncytium response properties depend on the rest period between runs. **a-d**, Astrocytes’ syncytium calcium response probability, onset, and strength in area M1/M2 depends on the rest period between runs. This inter-run interval dependency was accounted for during data analysis, including only trials with >15 s rest periods between runs. **a**, Paired traces of running activity (*left*) and syncytium calcium signals (*right*), ordered by the inter-trial interval and aligned on the current run’s onset (red lines). **b**, Astrocyte syncytium response probability as a function of the inter-run interval and trial type. **c**, Astrocyte syncytium response onset as a function of the inter-run interval and trial type. **d**, Astrocytes syncytium response strength, as quantified by response duration, peak, total activation extent, and mean pixel activation duration (*from left to right*), as a function of the inter-run interval and trial type.

**Extended Data Fig. 3.**
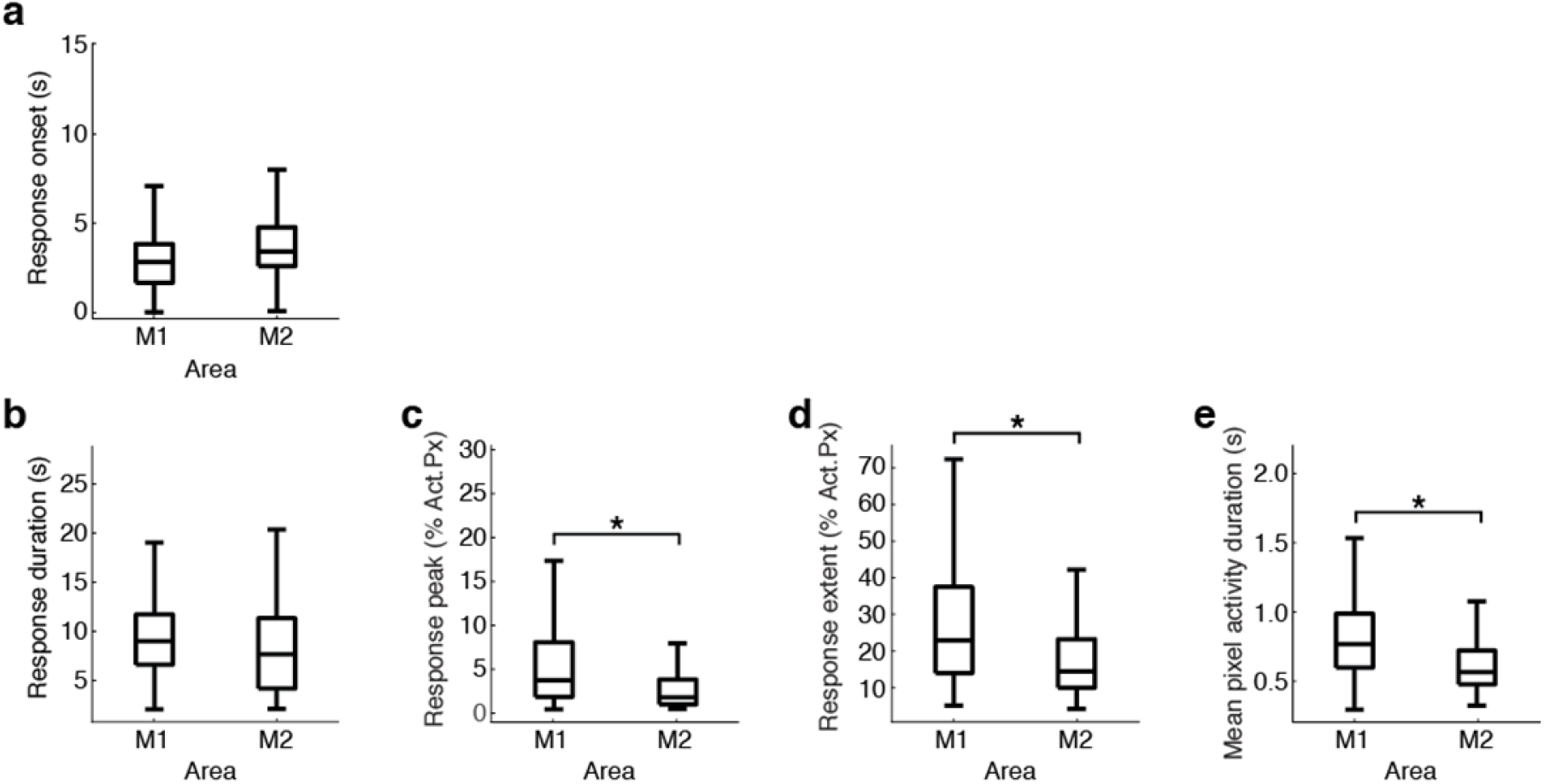
Astrocyte syncytium responses show regional differences. **a-e**, Astrocyte syncytium response onset and duration were comparable between areas M1 and M2 for the different trial types. In contrast, response peak, total activation extent, and mean pixel activation duration were significantly larger in area M1. **a**, Response onsets. **b**, Response durations. **c**, Response peaks. **d**, Total response extent. **e**, Mean pixel activation durations.

**Extended Data Fig. 4.**
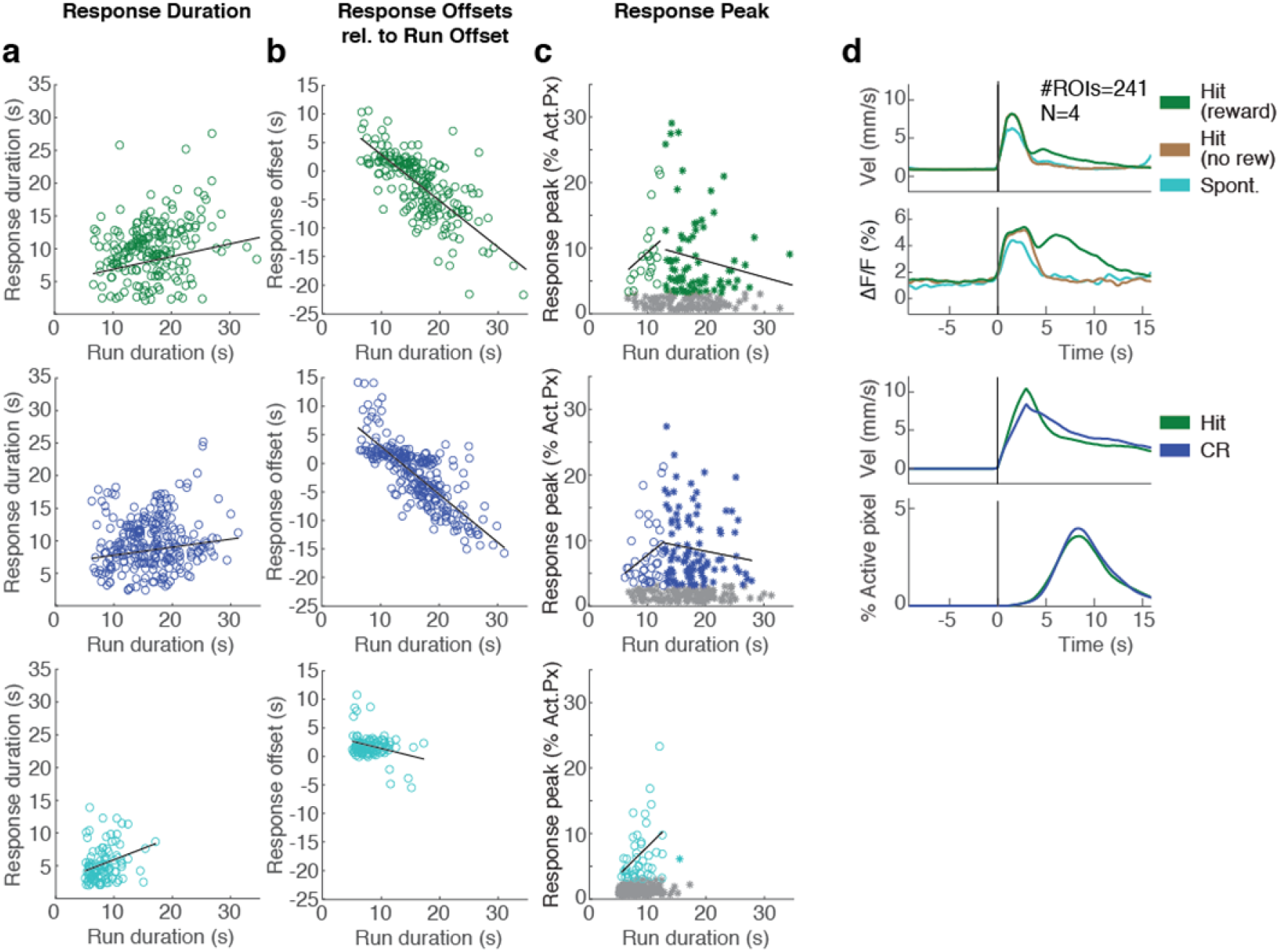
Astrocyte syncytium responses correlate with dopamine signaling in rewarded trials. **a**, Astrocyte syncytium responses in hit trials, CR trials, and spontaneous runs (*top to bottom*) increase slightly with run duration. **b**, Astrocyte syncytium response offsets relative to run offset decrease strongly for longer run durations in hit (*top*) and CR (*center*) trials. Response offsets coincide with run offsets in 13-15 s-long runs. In contrast, response offsets for spontaneous runs (*bottom*) are only slightly modulated by run duration, coinciding mostly with run offsets. **c**, Astrocyte syncytium response peak varies with run duration for hit trials, CR trials, and spontaneous runs (*top to bottom*). 13 s-long runs produced the highest response peaks. Linear fits to the data from <13 s and >13 s-long runs showed that response peak values steadily rose toward the “preferred” 13 s run duration and declined after that. Trials with response peaks of ≤3% active pixels (gray) were excluded from the linear fits. LME analysis was used to derive fit significance. **d**, Astrocyte response duration and offset in rewarded hit and CR trials correlated with the period dopamine is present in the extracellular space after reward delivery. Dopamine signals were measured with the genetically encoded indicator dLight1.2 in layer 2/3 of cortical areas M1/M2 during detection task performance^17^. *Top*, run velocity profiles and corresponding average dLight1.2 transients for rewarded hit trials (green), unrewarded hit trials (brown), and spontaneous runs (cyan) aligned at the run onset. The traces are an average across ROIs active during reward (241 ROIs from four mice). *Bottom*, run velocity profiles and corresponding astrocyte syncytium responses for rewarded hit (198 trials, green) and CR trials (260 trials, blue) aligned at the run onset.

**Extended Data Fig. 5.**
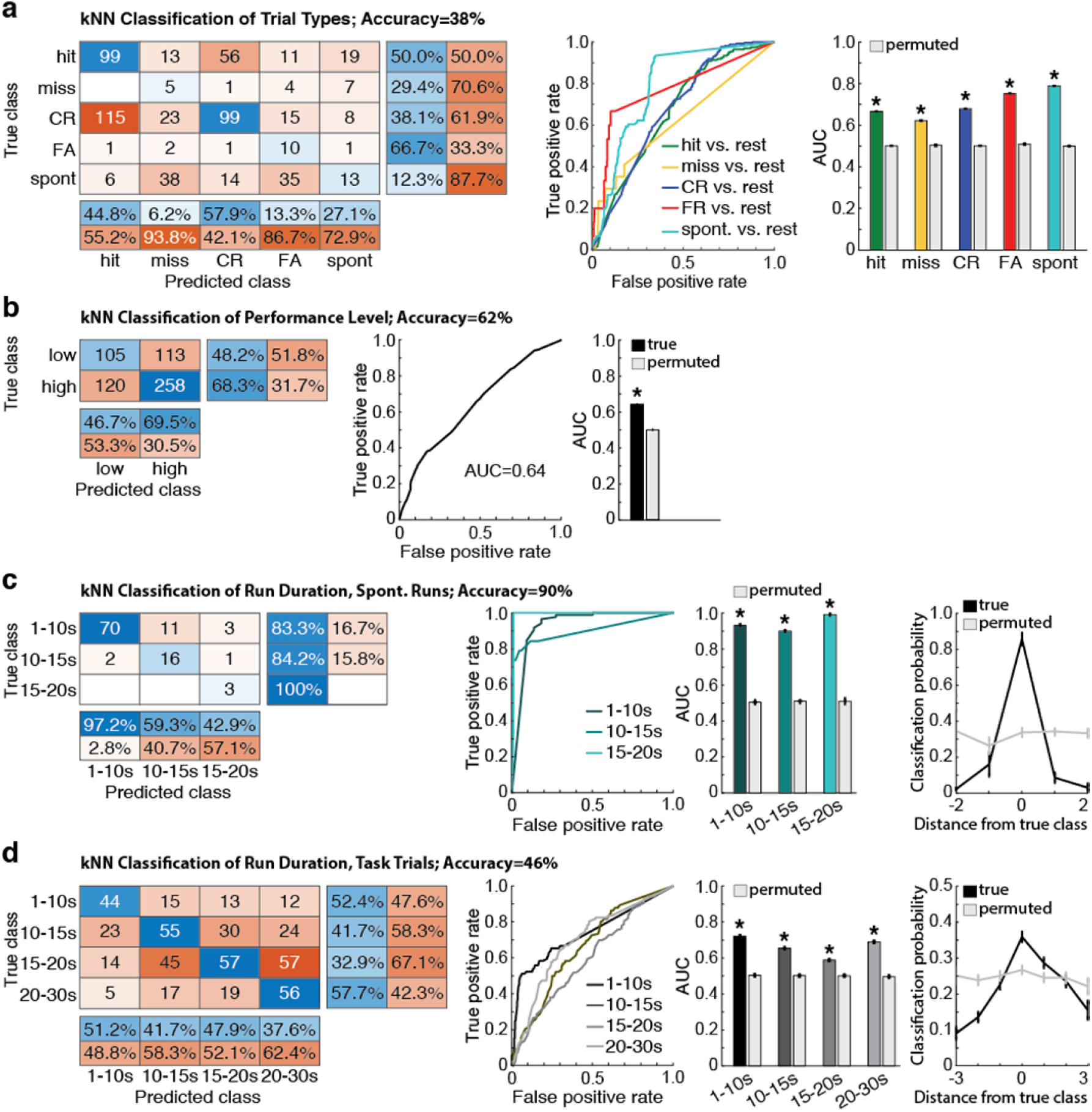
Behavioral aspects can be decoded from astrocytic syncytium responses using machine learning approaches. **a**-**d,** Astrocyte syncytium signals carry behavioral information as indicated by the k-nearest neighbor (kNN) classifier’s prediction accuracy. For each classification, confusion matrices, receiver-operating characteristic (ROC) curves and area under the ROC curves (AUC) for the classifier’s output, and statistical analysis of significance based on permutation tests are shown (*left to right*). Error bars indicate s.e.m. **a**, The kNN classifier decoded the trial type from astrocyte syncytium responses significantly above chance level. Most confusions happened between hit and CR trials. The decoding performance was worst for miss trials. **b**, Animal performance level could be significantly decoded from astrocyte syncytium signals. **c**, Spontaneous run durations could be decoded from astrocyte syncytium responses. **d**, Task-related run duration could be decoded from astrocyte syncytium responses. Confusions were more likely between neighboring run duration classes. Far-right plots in *c* and *d* show the decoding probabilities for a given run duration class as a function of distance from the true class (black line). The gray line depicts the average decoding probabilities based on permutation tests (see **Methods**).

**Extended Data Fig. 6.**
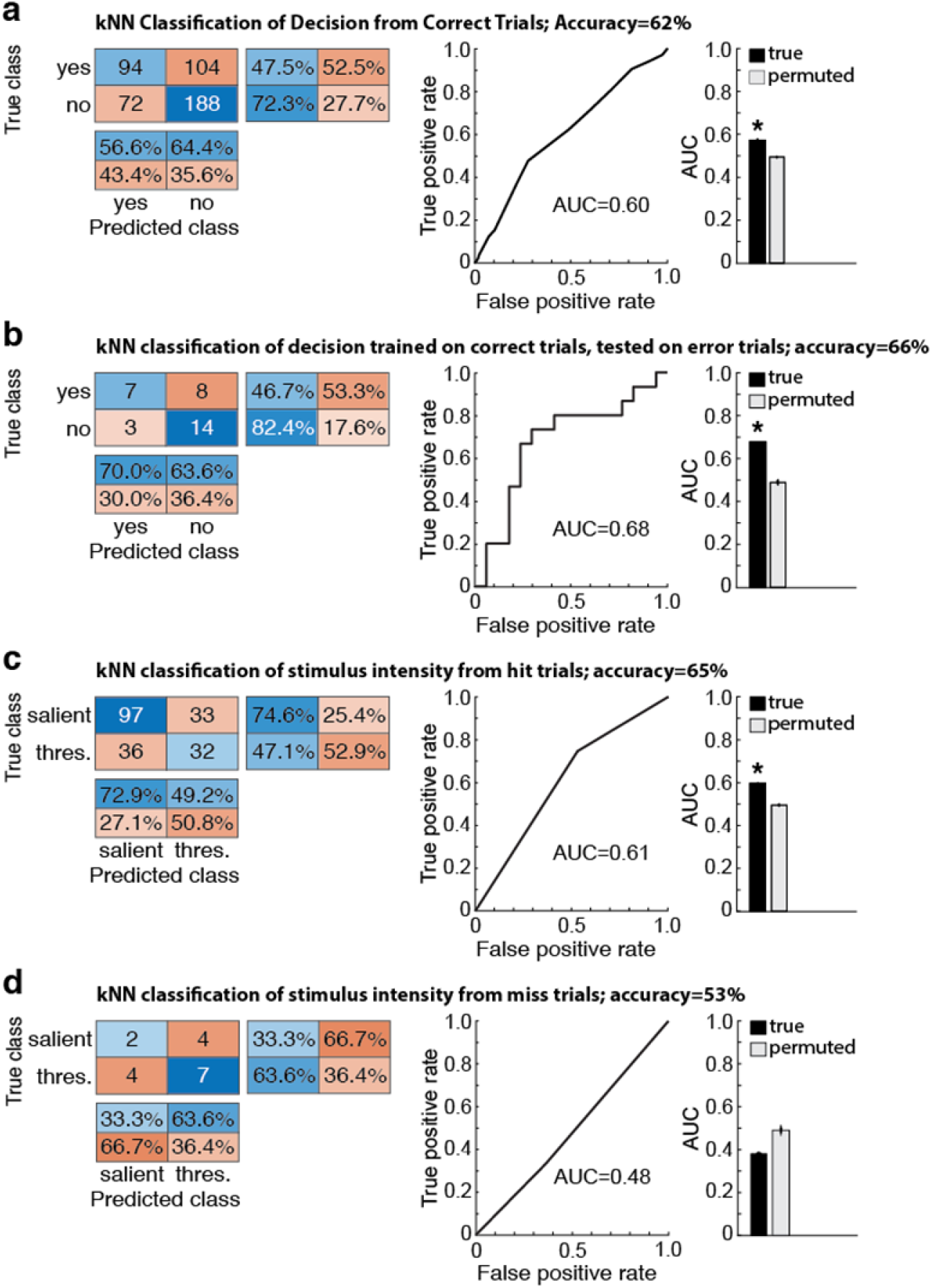
Astrocyte syncytium responses are behaviorally relevant. **a**-**d**, Separate classifications of correct and error trials reveal that information encoded by astrocyte syncytium calcium responses is relevant for the animal’s behavior. For each classification, confusion matrices, receiver-operating characteristic (ROC) curves for the classifier’s output and Area under the ROC curves (AUC), and statistical analysis of significance based on permutation tests are shown (*left to right*). Error bars indicate s.e.m. **a**, Decoding the animal’s decision about stimulus presence or absence was possible from astrocytes’ syncytium responses to hit and CR trials. **b**, Decoding the animal’s decision was also possible when the classifier was trained on correct (hit and CR) but tested on erroneous (miss and FA) trials. Significant prediction of the animal’s decision (miss-’no’, FA-’yes’) was confirmed by the AUC value. This value was significantly higher than AUC values obtained on a training set with randomly shuffled class labels. **c**, Information about stimulus intensity could be significantly decoded from astrocyte syncytium responses to hit trials. **d**, Miss trials lack behaviorally relevant sensory information, as the decoder fails to classify error trials according to stimulus intensity.

**Extended Data Fig. 7.**
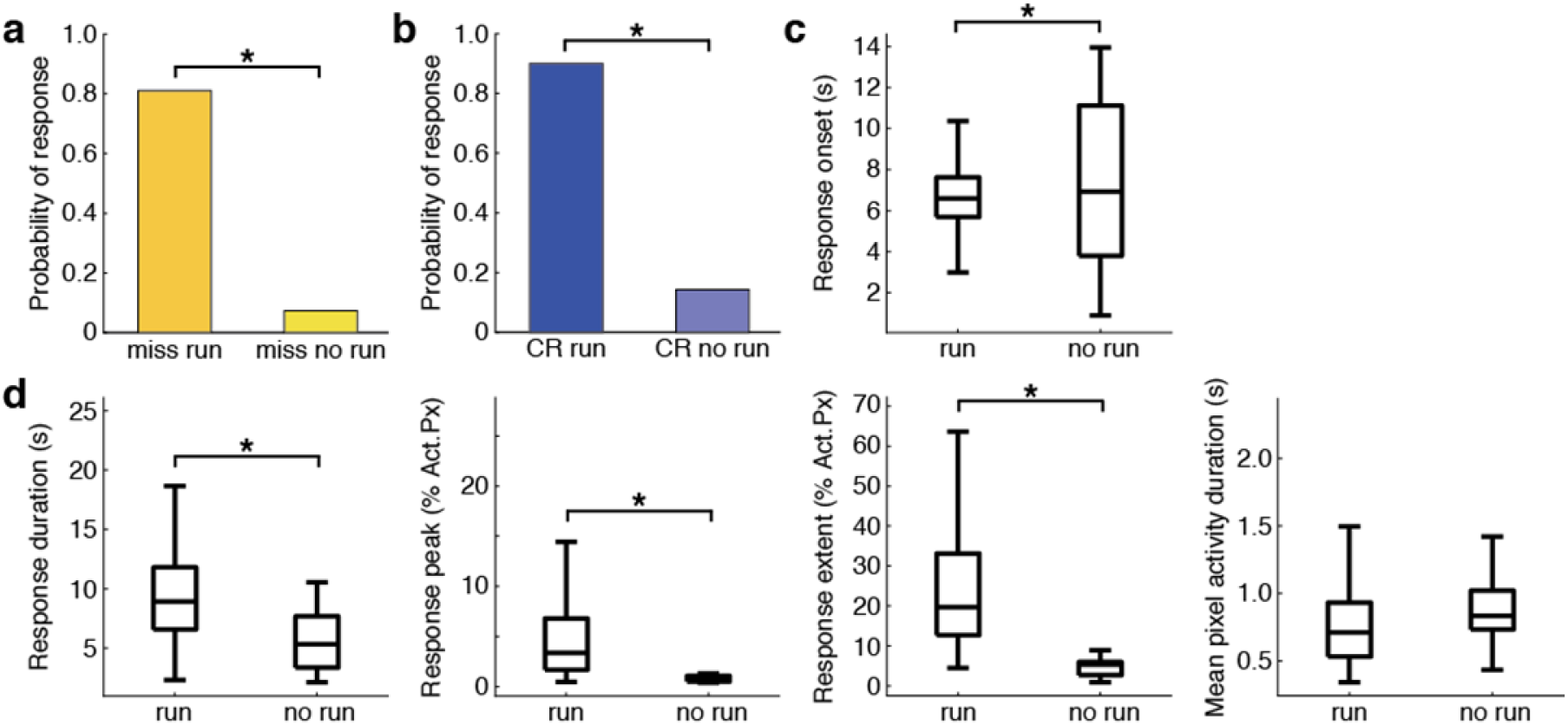
Astrocyte syncytium responses depend on run occurrence. **a**-**d**, Astrocyte syncytium responses are significantly different in miss and CR trials with or without a subsequent run. During miss and CR trials, the animals remain still throughout the stimulus presentation phase. However, they start running in most CR trials during reward consumption and occasionally miss trials after stimulus offset. These trial types allowed for the comparison of trials with and without a run. **a**, Astrocyte syncytium response probability for miss trials with and without a subsequent run. **b**, Astrocyte syncytium response probability for CR trials with and without a subsequent run. **c**, Response onsets for ‘run’ and ‘no run’ trials averaged across miss and CR trial types aligned at stimulus onset. **d**, Astrocyte syncytium response strength, as quantified by response duration, peak, total activation extent, and mean pixel activation (*from left to right*), for ‘run’ and ‘no run’ trials averaged across miss and CR trial types.

## Extended Data Tables

**Table 1.** Numbers of trials included in the syncytium response analysis.

|  | Hit | Miss |  | CR |  | FA | Spont. |
| --- | --- | --- | --- | --- | --- | --- | --- |
|  |  | run | no run | run | no run |  |  |
| **Significant response** | 198 | 17 | 253 | 260 | 6 | 15 | 106 |
| **No significant response** | 23 | 4 | 20 | 30 | 1 | 3 | 61 |

**Table 2.** Generalized linear mixed-effects model for astrocyte syncytium response probability. All qualifying trials (Table 1), followed by a run, were included in the analysis. Degrees of freedom: 717.

| **Behavioral Variable** | **Coefficient** | **Standard Error** | **p-value** |
| --- | --- | --- | --- |
| Trial type hit vs. miss | 3.48 | 2.93 | 0.23 |
| Trial type hit vs. CR | -0.51 | 1.03 | 0.62 |
| Trial type hit vs. FA | -2.94 | 3.61 | 0.42 |
| Trial type hit vs. spont. | -2.96 | 1.12 | 0.01 |
| Trial type miss vs. CR | 0.66 | 0.64 | 0.30 |
| Trial type miss vs. FA | -0.01 | 0.88 | 0.99 |
| Trial type miss vs. spont. | -0.83 | 0.61 | 0.18 |
| Trial type CR vs. FA | -0.68 | 0.70 | 0.33 |
| Trial type CR vs. spont. | -1.49 | 0.29 | $5 \cdot 10^{-7}$ |
| Trial type FA vs. spont. | -0.81 | 0.68 | 0.23 |
| Run duration | 0.12 | 0.05 | 0.02 |
| Inter-run interval | 0.04 | 0.01 | $5 \cdot 10^{-4}$ |
| Trial type miss: Run duration | -0.44 | 0.34 | 0.19 |
| Trial type CR: Run duration | 0.02 | 0.07 | 0.74 |
| Trial type FA: Run duration | 0.43 | 0.51 | 0.40 |
| Trial type spont.: Run duration | 0.31 | 0.12 | 0.01 |

**Table 3.** Linear mixed-effects model for astrocyte syncytium response onset. All qualifying trials (Table 1) with a run and significant syncytium response were included. Degrees of freedom: 590.

| Behavioral Variable | Coefficient | Standard Error | p-value |
| --- | --- | --- | --- |
| Trial type hit vs. miss | -0.07 | 0.40 | 0.85 |
| Trial type hit vs. CR | 0.81 | 0.15 | $1 \cdot 10^{-7}$ |
| Trial type hit vs. FA | 0.64 | 0.41 | 0.12 |
| Trial type hit vs. spont. | 0.60 | 0.20 | 0.002 |
| Trial type miss vs. CR | 0.88 | 0.40 | 0.03 |
| Trial type miss vs. FA | 0.71 | 0.54 | 0.19 |
| Trial type miss vs. spont. | 0.68 | 0.40 | 0.09 |
| Trial type CR vs. FA | -0.17 | 0.41 | 0.67 |
| Trial type CR vs. spont. | -0.21 | 0.19 | 0.29 |
| Trial type FA vs. spont. | 0.03 | 0.42 | 0.94 |
| Inter-run interval | 0.04 | 0.01 | $5 \cdot 10^{-4}$ |

**Table 4.** Linear mixed-effects model for astrocyte syncytium response duration. All qualifying trials (Table 1) with a run and significant syncytium response were included. Degrees of freedom: 589.

| Behavioral Variable | Coefficient | Standard Error | p-value |
| --- | --- | --- | --- |
| Trial type hit vs. miss | -2.04 | 0.91 | 0.02 |
| Trial type hit vs. CR | -0.63 | 0.33 | 0.06 |
| Trial type hit vs. FA | -3.82 | 0.94 | $5 \cdot 10^{-5}$ |
| Trial type hit vs. spont. | -3.16 | 0.48 | $1 \cdot 10^{-10}$ |
| Trial type miss vs. CR | 1.40 | 0.90 | 0.12 |
| Trial type miss vs. FA | -1.78 | 1.19 | 0.13 |
| Trial type miss vs. spont. | -1.12 | 0.88 | 0.20 |
| Trial type CR vs. FA | -3.19 | 0.93 | $6 \cdot 10^{-4}$ |
| Trial type CR vs. spont. | -2.52 | 0.47 | $9 \cdot 10^{-8}$ |
| Trial type FA vs. spont. | -0.66 | 0.92 | 0.47 |
| Run duration | 0.17 | 0.03 | $3 \cdot 10^{-7}$ |
| Inter-run interval | 0.03 | 0.01 | $1 \cdot 10^{-6}$ |

**Table 5.** Linear mixed-effects model for astrocyte syncytium response peak. All qualifying trials (Table 1) with a run and significant syncytium response were included. Degrees of freedom: 588.

| Behavioral Variable | Coefficient | Standard Error | p-value |
| --- | --- | --- | --- |
| Trial type hit vs. miss | 0.04 | 0.22 | 0.87 |
| Trial type hit vs. CR | -0.08 | 0.08 | 0.31 |
| Trial type hit vs. FA | -0.61 | 0.22 | $6 \cdot 10^{-3}$ |
| Trial type hit vs. spont. | -0.33 | 0.10 | $2 \cdot 10^{-3}$ |
| Trial type miss vs. CR | -0.12 | 0.21 | 0.58 |
| Trial type miss vs. FA | -0.65 | 0.3 | 0.03 |
| Trial type miss vs. spont. | -0.36 | 0.22 | 0.09 |
| Trial type CR vs. FA | -0.53 | 0.22 | 0.02 |
| Trial type CR vs. spont. | -0.24 | 0.10 | 0.02 |
| Trial type FA vs. spont. | 0.28 | 0.22 | 0.20 |
| Area (M2) | -0.48 | 0.19 | $1 \cdot 10^{-83}$ |
| Performance (high) | -0.29 | 0.08 | $1 \cdot 10^{-4}$ |
| Inter-run interval | 0.01 | 0.001 | $7 \cdot 10^{-13}$ |

**Table 6.** Linear mixed-effects model for astrocyte syncytium response extent. All qualifying trials (Table 1) with a run and significant syncytium response were included. Degrees of freedom: 588.

| Behavioral Variable | Coefficient | Standard Error | p-value |
| --- | --- | --- | --- |
| Trial type hit vs. miss | -0.31 | 0.14 | 0.03 |
| Trial type hit vs. CR | -0.12 | 0.05 | 0.03 |
| Trial type hit vs. FA | -0.64 | 0.15 | $2 \cdot 10^{-5}$ |
| Trial type hit vs. spont. | -0.54 | 0.07 | $3 \cdot 10^{-14}$ |
| Trial type miss vs. CR | 0.19 | 0.14 | 0.18 |
| Trial type miss vs. FA | -0.33 | 0.19 | 0.08 |
| Trial type miss vs. spont. | -0.23 | 0.14 | 0.10 |
| Trial type CR vs. FA | -0.52 | 0.14 | $4 \cdot 10^{-4}$ |
| Trial type CR vs. spont. | -0.42 | 0.08 | $1 \cdot 10^{-9}$ |
| Trial type FA vs. spont. | 0.10 | 0.15 | 0.50 |
| Area (M2) | -0.26 | 0.12 | 0.03 |
| Performance (high) | -0.18 | 0.05 | $2 \cdot 10^{-4}$ |
| Inter-run interval | 0.008 | 0.001 | $6 \cdot 10^{-14}$ |

**Table 7.** Linear mixed-effects model for mean pixel activity duration. All qualifying trials (Table 1) with a run and significant syncytium response were included. Degrees of freedom: 588.

| Behavioral Variable | Coefficient | Standard Error | p-value |
| --- | --- | --- | --- |
| Trial type hit vs. miss | -0.09 | 0.07 | 0.23 |
| Trial type hit vs. CR | -0.05 | 0.03 | 0.60 |
| Trial type hit vs. FA | -0.25 | 0.07 | $7 \cdot 10^{-4}$ |
| Trial type hit vs. spont. | -0.17 | 0.03 | $5 \cdot 10^{-7}$ |
| Trial type miss vs. CR | 0.03 | 0.07 | 0.62 |
| Trial type miss vs. FA | -0.16 | 0.09 | 0.09 |
| Trial type miss vs. spont. | -0.09 | 0.07 | 0.21 |
| Trial type CR vs. FA | -0.19 | 0.07 | 0.006 |
| Trial type CR vs. spont. | -0.12 | 0.03 | $2 \cdot 10^{-4}$ |
| Trial type FA vs. spont. | 0.07 | 0.07 | 0.31 |
| Area (M2) | -0.20 | 0.09 | 0.02 |
| Performance (high) | -0.07 | 0.02 | 0.005 |
| Inter-run interval | 0.003 | $5 \cdot 10^{-4}$ | $3 \cdot 10^{-10}$ |

**Table 8.** Linear mixed-effects models for astrocyte syncytium response strength probing correct/error trial encoding. The categorical variable correct/error was included instead of trial type. All qualifying task trials (Table 1) with a run and significant syncytium response were included. Degrees of freedom: 486.

| Behavioral Variable | Coefficient | Standard Error | p-value |
| --- | --- | --- | --- |
| **Response duration** |  |  |  |
| Correct/Error trials | -2.66 | 0.73 | $3 \cdot 10^{-4}$ |
| Run duration | 0.16 | 0.04 | $5 \cdot 10^{-6}$ |
| Inter-run interval | 0.03 | 0.007 | $6 \cdot 10^{-6}$ |
| **Extent** |  |  |  |
| Correct/Error trials | -0.40 | 0.10 | $2 \cdot 10^{-4}$ |
| Performance (high) | -0.15 | 0.05 | 0.005 |
| Inter-run interval | 0.01 | 0.001 | $2 \cdot 10^{-12}$ |
| **Mean pixel activity duration** |  |  |  |
| Correct/Error trials | -0.11 | 0.05 | 0.03 |
| Area (M2) | -0.21 | 0.09 | 0.03 |
| Inter-run interval | 0.003 | $5 \cdot 10^{-4}$ | $2 \cdot 10^{-9}$ |

**Table 9.**
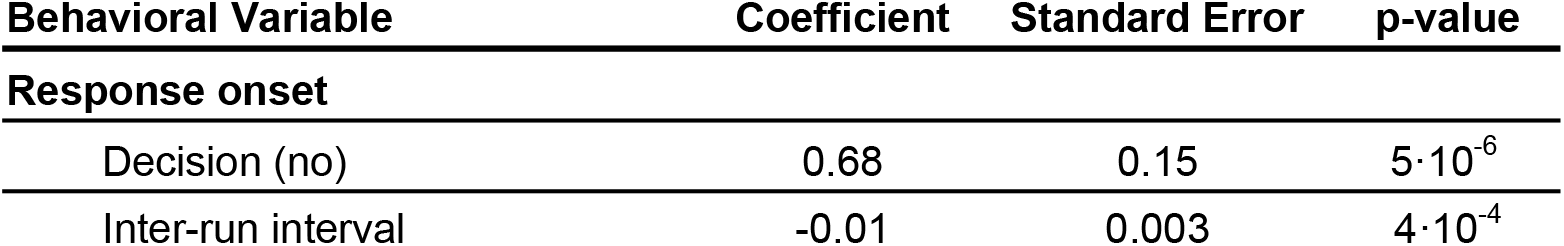
Linear mixed-effects model for astrocyte syncytium response onset probing decision encoding (‘yes’ - hit and FA trials; ‘no’ - CR and miss trials). The categorical variable decision was included instead of the trial type. All qualifying task trials (Table 1) with a run and significant syncytium response were included. Degrees of freedom: 487.

**Table 10.**
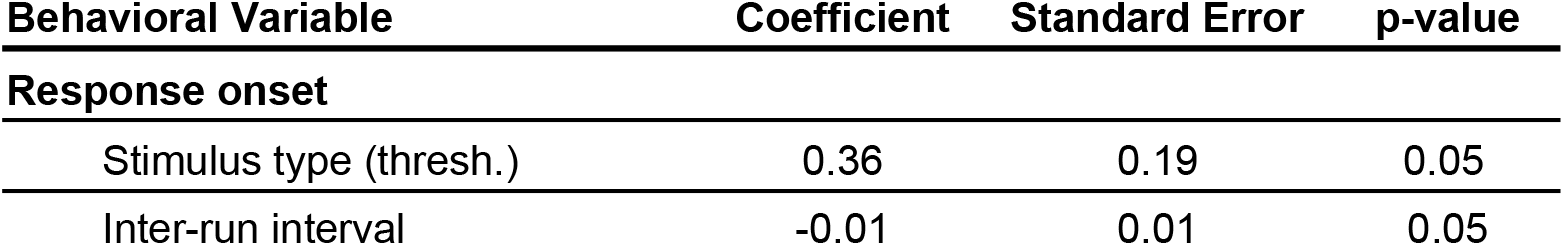
Linear mixed-effects model testing the relationship between astrocyte syncytium response and stimulus intensity. Only hit trials with a run and a significant astrocyte syncytium response were included in the analysis (Table 1). Degrees of freedom: 240.

**Table 11.** Linear mixed-effects models examining the astrocyte syncytium response’s probability, onset, and strength with respect to run occurrence. All trials with a significant syncytium response were included (Table 1). All trials were aligned at trial onset.

| Behavioral Variable | Coefficient | Standard Error | p-value | Degrees of Freedom |
| --- | --- | --- | --- | --- |
| **Probability (miss trials)** |  |  |  |  |
| Run/No run (run) | 4.2 | 0.74 | $1 \cdot 10^{-12}$ | 292 |
| **Probability (CR trials)** |  |  |  |  |
| Run/No run (run) | 4.93 | 1.32 | $2 \cdot 10^{-4}$ | 295 |
| **Onset** |  |  |  |  |
| Run/No run (run) | -2.10 | 0.43 | $2 \cdot 10^{-5}$ | 297 |
| **Response duration** |  |  |  |  |
| Run/No run (run) | 3.50 | 0.86 | $6 \cdot 10^{-5}$ | 294 |
| **Response peak** |  |  |  |  |
| Run/No run (run) | 1.45 | 0.22 | $9 \cdot 10^{-11}$ | 294 |
| **Response extent** |  |  |  |  |
| Run/No run (run) | 1.75 | 0.14 | $2 \cdot 10^{-29}$ | 297 |
| **Mean pixel activity duration** |  |  |  |  |
| Run/No run (run) | -0.09 | 0.07 | 0.25 | 297 |

**Table 12.** Linear mixed-effects model for astrocyte syncytium response duration, testing for the effect of run duration in hit trials, CR trials, and spontaneous runs. All qualifying trials (Table 1) with a run and significant syncytium response for the respective trial types were included.

| Behavioral Variable | Coefficient | Standard Error | p-value | Degrees of Freedom |
| --- | --- | --- | --- | --- |
| **Hit trials** |  |  |  |  |
| Run duration | 0.19 | 0.05 | $4 \cdot 10^{-4}$ | 195 |
| Inter-run interval | 0.06 | 0.01 | $9 \cdot 10^{-5}$ | 195 |
| **CR trials** |  |  |  |  |
| Run duration | 0.12 | 0.05 | 0.01 | 257 |
| Inter-run interval | 0.02 | 0.01 | 0.003 | 257 |
| **Spontaneous runs** |  |  |  |  |
| Run duration | 0.34 | 0.10 | $6 \cdot 10^{-4}$ | 104 |

**Table 13.**
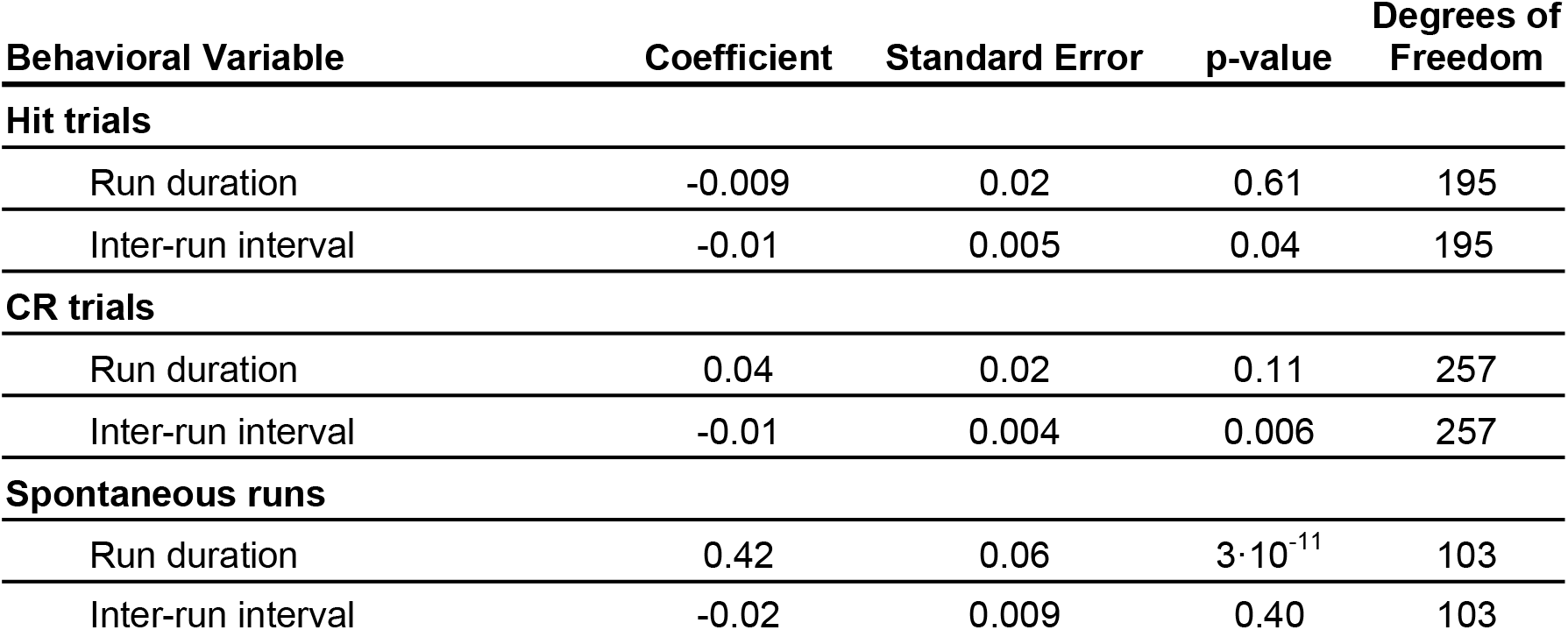
Linear mixed-effects model for astrocyte syncytium response onset, testing for the effect of run duration in hit trials, CR trials, and spontaneous runs. All qualifying trials (Table 1) with a run and significant syncytium response for the respective trial types were included.

| Behavioral Variable | Coefficient | Standard Error | p-value | Degrees of Freedom |
| --- | --- | --- | --- | --- |
| **Hit trials** |  |  |  |  |
| Run duration | -0.009 | 0.02 | 0.61 | 195 |
| Inter-run interval | -0.01 | 0.005 | 0.04 | 195 |
| **CR trials** |  |  |  |  |
| Run duration | 0.04 | 0.02 | 0.11 | 257 |
| Inter-run interval | -0.01 | 0.004 | 0.006 | 257 |
| **Spontaneous runs** |  |  |  |  |
| Run duration | 0.42 | 0.06 | $3 \cdot 10^{-11}$ | 103 |
| Inter-run interval | -0.02 | 0.009 | 0.40 | 103 |

**Table 14.**
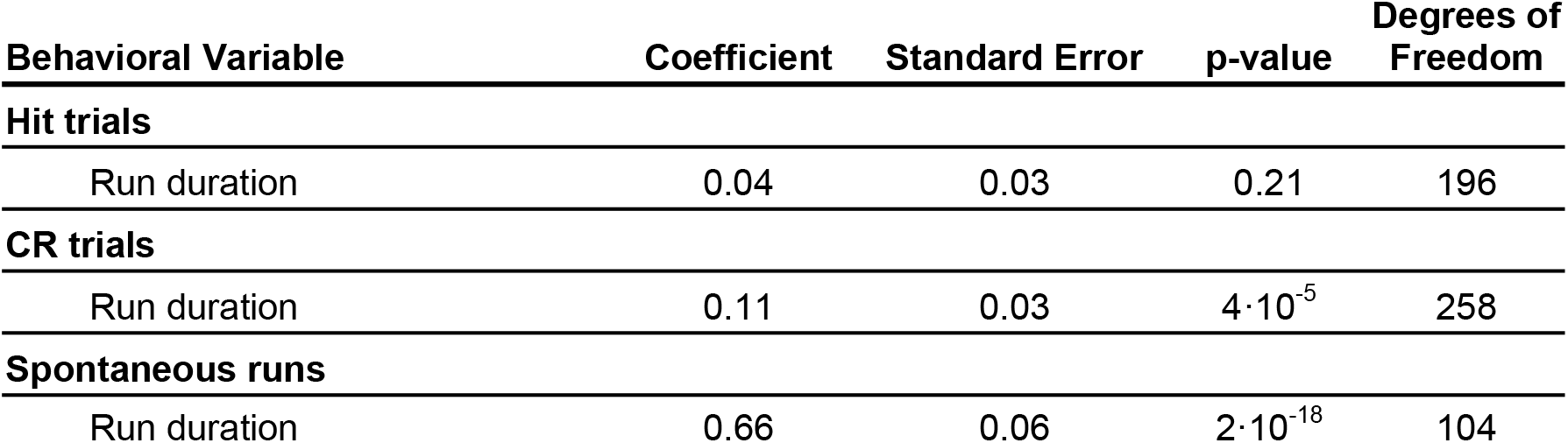
Linear mixed-effects model for the astrocyte syncytium response’s peak location testing for the effect of run duration in hit trials, CR trials, and spontaneous runs. All qualifying trials (Table 1) with a run and significant syncytium response for the respective trial type were included.

| Behavioral Variable | Coefficient | Standard Error | p-value | Degrees of Freedom |
| --- | --- | --- | --- | --- |
| **Hit trials** |  |  |  |  |
| Run duration | 0.04 | 0.03 | 0.21 | 196 |
| **CR trials** |  |  |  |  |
| Run duration | 0.11 | 0.03 | $4 \cdot 10^{-5}$ | 258 |
| **Spontaneous runs** |  |  |  |  |
| Run duration | 0.66 | 0.06 | $2 \cdot 10^{-18}$ | 104 |

**Table 15.** Linear mixed-effects model for astrocyte syncytium response offset aligned on run onset testing for the effect of run duration in hit trials, CR trials, and spontaneous runs. All qualifying trials (Table 1) with a run and significant syncytium response for the respective trial types were included.

| Behavioral Variable | Coefficient | Standard Error | p-value | Degrees of Freedom |
| --- | --- | --- | --- | --- |
| **Hit trials** |  |  |  |  |
| Run duration | 0.18 | 0.05 | $3 \cdot 10^{-4}$ | 195 |
| Inter-run interval | 0.50 | 0.01 | $4 \cdot 10^{-4}$ | 195 |
| **CR trials** |  |  |  |  |
| Run duration | 0.16 | 0.04 | $6 \cdot 10^{-5}$ | 257 |
| Inter-run interval | 0.01 | 0.006 | 0.05 | 257 |
| **Spontaneous runs** |  |  |  |  |
| Run duration | 0.75 | 0.08 | $8 \cdot 10^{-16}$ | 104 |

**Table 16.** Linear mixed-effects model for astrocyte syncytium response offset aligned at the run offset, testing for the effect of run duration in hit trials, CR trials, and spontaneous runs. All qualifying trials (Table 1) with a run and significant syncytium response for the respective trial types were included.

| Behavioral Variable | Coefficient | Standard Error | p-value | Degrees of Freedom |
| --- | --- | --- | --- | --- |
| **Hit trials** |  |  |  |  |
| Run duration | -0.82 | 0.05 | $2 \cdot 10^{-37}$ | 195 |
| Inter-run interval | 0.50 | 0.10 | $5 \cdot 10^{-4}$ | 195 |
| **CR trials** |  |  |  |  |
| Run duration | -0.84 | 0.04 | $2 \cdot 10^{-59}$ | 257 |
| Inter-run interval | 0.01 | 0.006 | 0.05 | 257 |
| **Spontaneous runs** |  |  |  |  |
| Run duration | -0.25 | 0.08 | 0.002 | 104 |

**Table 17.**
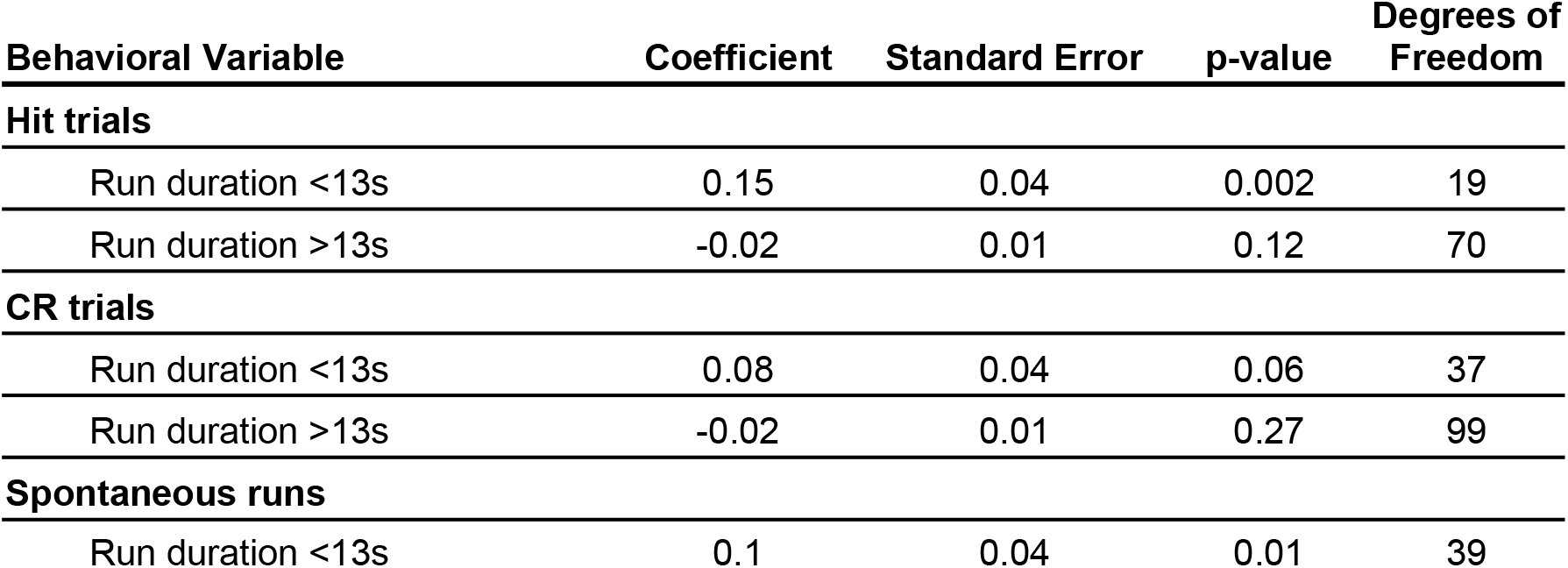
Linear mixed-effects model for determining the relationship between syncytium response peaks and run duration. All qualifying trials (Table 1) with a run and significant syncytium response (>3% active pixels peak value) were included.

**Table 18.** Optimized hyperparameters used in the kNN classification analyses.

| Classification | Nr. neighbors | Distance | Distance weight |
| --- | --- | --- | --- |
| Rewarded trials/ spontaneous runs | 7 | correlation | equal |
| Rewarded/ error, non-rewarded | 7 | correlation | equal |
| Trial types (hit, miss, CR, FA, spont.) | 9 | correlation | equal |
| Performance | 11 | jaccard | equal |
| Run duration | 5 | cosine | equal |
| Decision | 50 | correlation | inverse |

correlation: linear correlation between observations (data were treated as a sequence of values) jaccard: Jaccard coefficient (the percentage of nonzero coordinates that differ)
cosine: cosine of the angle between observations (data were treated as vectors)

**Table 19.**
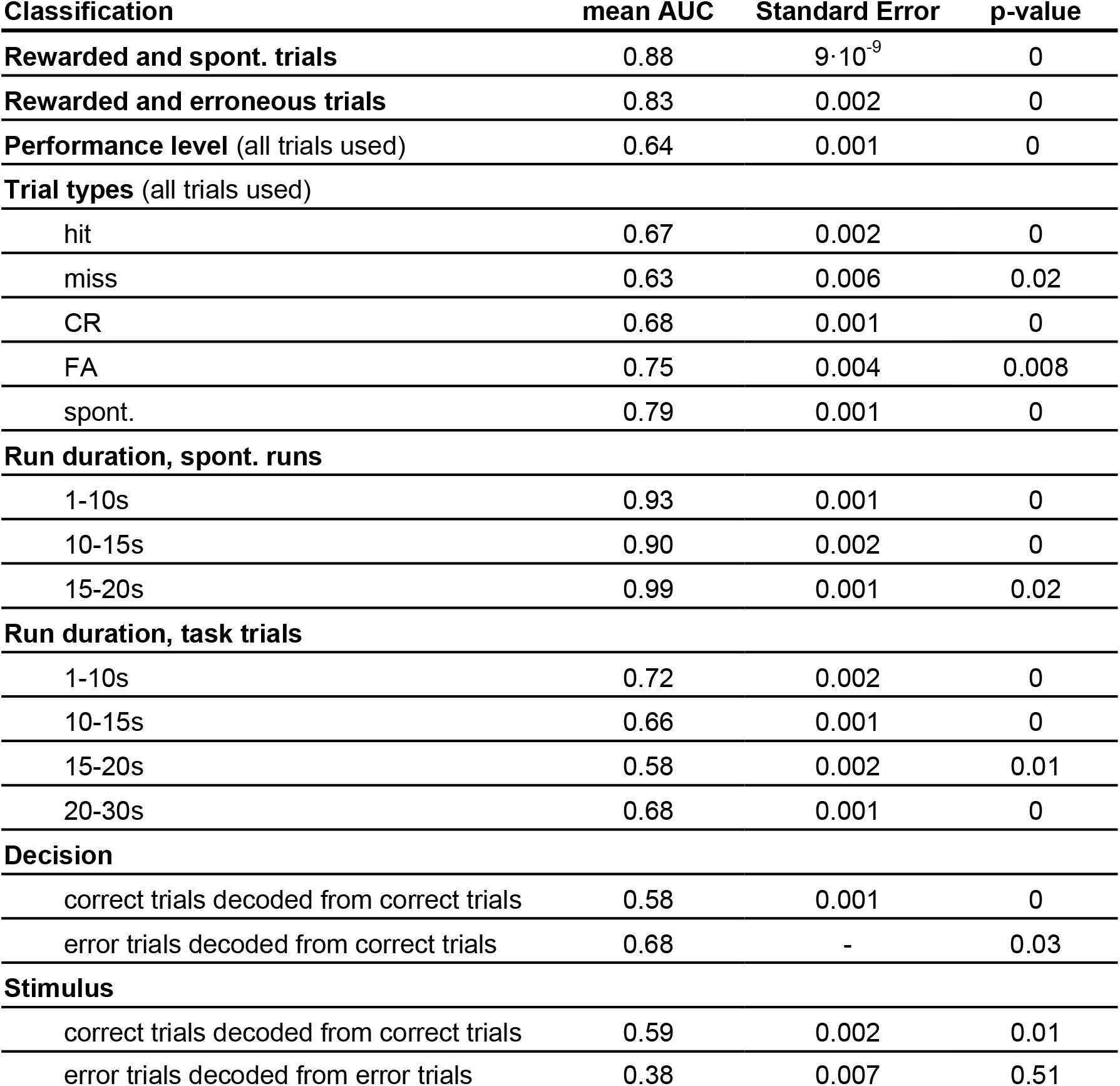
Permutation tests evaluating the kNN classifier’s decoding performance. All qualifying trials (Table 1) with a run and significant syncytium response for the respective trial types were included.

| Classification | mean AUC | Standard Error | p-value |
| --- | --- | --- | --- |
| **Rewarded and spont. trials** | 0.88 | $9 \cdot 10^{-9}$ | 0 |
| **Rewarded and erroneous trials** | 0.83 | 0.002 | 0 |
| **Performance level** (all trials used) | 0.64 | 0.001 | 0 |
| **Trial types** (all trials used) |  |  |  |
| hit | 0.67 | 0.002 | 0 |
| miss | 0.63 | 0.006 | 0.02 |
| CR | 0.68 | 0.001 | 0 |
| FA | 0.75 | 0.004 | 0.008 |
| spont. | 0.79 | 0.001 | 0 |
| **Run duration, spont. runs** |  |  |  |
| 1-10s | 0.93 | 0.001 | 0 |
| 10-15s | 0.90 | 0.002 | 0 |
| 15-20s | 0.99 | 0.001 | 0.02 |
| **Run duration, task trials** |  |  |  |
| 1-10s | 0.72 | 0.002 | 0 |
| 10-15s | 0.66 | 0.001 | 0 |
| 15-20s | 0.58 | 0.002 | 0.01 |
| 20-30s | 0.68 | 0.001 | 0 |
| **Decision** |  |  |  |
| correct trials decoded from correct trials | 0.58 | 0.001 | 0 |
| error trials decoded from correct trials | 0.68 | - | 0.03 |
| **Stimulus** |  |  |  |
| correct trials decoded from correct trials | 0.59 | 0.002 | 0.01 |
| error trials decoded from error trials | 0.38 | 0.007 | 0.51 |

